# Genome assembly and analysis of *Lactuca virosa*: implications for lettuce breeding

**DOI:** 10.1101/2023.05.03.539295

**Authors:** Wei Xiong, Dirk-Jan M. van Workum, Lidija Berke, Linda V. Bakker, Elio Schijlen, Frank F.M. Becker, Henri van de Geest, Sander Peters, Richard Michelmore, Rob van Treuren, Marieke Jeuken, Sandra Smit, M. Eric Schranz

## Abstract

Lettuce (*Lactuca sativa* L.) is a leafy vegetable crop with ongoing breeding efforts related to quality, resilience, and innovative production systems. Genetic variation of important traits in close relatives is necessary to meet lettuce breeding goals. *Lactuca virosa* (2x=2n=18), a wild relative assigned to the tertiary lettuce gene pool, has a much larger genome (3.7 Gbp) than *Lactuca sativa* (2.5 Gbp). It has been used in interspecific crosses and is a donor to modern crisphead lettuce cultivars. Here, we present a *de novo* reference assembly of *L. virosa* with high continuity and complete gene space. This assembly facilitated comparisons to the genome of *L. sativa* and to that of the wild species *L. saligna*, a representative of the secondary lettuce gene pool. To assess the diversity in gene content, we classified the genes of the three *Lactuca* species as core, accessory and unique. In addition, we identified three interspecific chromosomal inversions compared to *L. sativa*, which each may cause recombination suppression and thus hamper future introgression breeding. Using three-way comparisons in both reference-based and reference-free manners, we show that the proliferation of long-terminal repeat elements has driven the genome expansion of *L. virosa*. Further, we performed a genome-wide comparison of immune genes, nucleotide-binding leucine-rich repeat, and receptor-like kinases among *Lactuca* spp. and indicate the evolutionary patterns and mechanisms behind their expansions. These genome analyses greatly facilitate the understanding of genetic variation in *L. virosa*, which is beneficial for the breeding of improved lettuce varieties.

## 1. Introduction

Lettuce (*Lactuca sativa* L.) is a crop with an economic value of ∼3 billion USD per year [1]. To develop better lettuce cultivars, breeders often search for novel genetic variations in lettuce wild relatives. *Lactuca virosa* (biennial) is a donor for resistance to different pests and pathogens and a representative species in the lettuce gene pool [2–5]. The exploitation of *L. virosa* for lettuce breeding has had both challenges and successes. For example, despite reproductive barriers for direct intercrossing with lettuce, breeders and scientists were able to execute interspecific hybridization bridged by *L. serriola* to introduce traits like robust root architecture and resistance to currant-lettuce aphid, downy mildew and viruses [6–8]. Such interspecific crosses are part of the breeding pedigrees of the well-known cultivars Vanguard and Salinas, representing modern crisphead lettuce cultivars [9,10]. Novel introgressions of desired genes and traits from *L. virosa* into cultivated lettuce could be realized through an improved understanding of its genomic content.

A reference genome and derived molecular markers are essential for breeders to select traits accurately and trace introgressions in cultivated lettuce from *L. virosa*. For example, genome-wide association studies (GWAS) have been performed to identify SNP variants [10] that are associated with interesting traits in lettuce [11–13] using the assembled lettuce (*L. sativa*) reference genome [14], which can be used to develop markers for lettuce breeding to accelerate selection in offspring [15]. In addition to GWAS, a reference genome of *L. virosa* will also facilitate genomic analyses for various biological questions. A whole-genome screening can search genetic determinants, for instance, that trigger a resistance response. Genome rearrangements can be detected between *L. virosa* and other *Lactuca* species via comparative genomics.

The creation of a reference genome for *L. virosa* is challenging, because, even though it is a diploid species (2n=2x=18), it has a considerable larger genome (3.7 Gbp) than *L. sativa* (2.5 Gbp)and *L. saligna* (2.3 Gbp) [16]. This is likely due to transposable elements [17]. To date, there is only a single available genome assembly of *L. virosa* (CGN04683) [18], which is a short-read based and highly fragmented assembly (3,694,810 scaffolds; N50=4,910bp) with a relatively high completeness (BUSCO=92.7%). Long-read sequencing could significantly improve the accuracy and continuity of a *L. virosa* genome assembly.

Here, we present a near chromosome-level *de novo* assembly of *L. virosa* (CGN04683) using a combination of long-read and short-read sequencing plus Bionano and Dovetail scaffolding. We contextualize the *L. virosa* genome within the lettuce gene pool together with the *L. sativa* and *L. saligna* [19] genomes. First, we show shared and specific homology groups across the three species. Based on homologs, we show interspecific collinearity with an emphasis on inversions in different chromosomes. Next, we demonstrate that the proliferation of long-terminal repeat (LTR) superfamilies underlies the genome expansion of *L. virosa*. Finally, we describe a well-classified inventory of the two important resistance-related gene types encoding nucleotide-binding leucine-rich repeat (NLR) receptors and receptor-like kinases (RLK).

## 2. Materials and Methods

### 2.1 DNA and RNA sequencing

*L. virosa* accession CGN04683, is also known as IVT280 and is resistant to *Nasonovia ribisnigri* (currant-lettuce aphid) [7]. Single seed descent of accession ‘IVT280’ (seeds obtained from a breeding company) was grown for whole-genome sequencing. The seeds were stratified at 4°C for three days to improve germination. Subsequently, seedlings were grown in a growth chamber at 18–21°C and a relative humidity of 75–78%. After eight weeks, plants were transplanted to larger pots containing potting soil and grown under greenhouse conditions. Tissue sampling was performed when plants were close to bolting, and DNA was extracted using the same protocol described in Xiong *et al*. [18]. DNA material was used to prepare appropriate libraries and to produce a 20-fold coverage of long-read data generated by PacBio Sequel technology and a 69-fold coverage of paired-end (PE) reads obtained by Illumina sequencing. An optical mapping library of 130X coverage was produced by Bionano mapping for hybrid scaffolding. A Hi-C library produced by Dovetail Genomics provided 10,492X physical coverage of the genome (10 kbp – 10 Mbp pairs) for *in vitro* proximity ligation (Supplementary Table 1). As additional evidence for gene prediction, RNA was isolated from pooled samples of leaf, root, and flower tissues (pooled from different floral stages) using a Direct Zol RNA Miniprep Plus kit (Zymo Research) followed by treatment with DNAse. RNA was purified by ethanol precipitation. The concentration and purity of RNA samples were measured with a Nanodrop 2000c spectrophotometer and a Qubit 4.0 fluorometer using an RNA Broad Range assay (Thermo Fisher Scientific). PE sequencing (2 x 125 bp) was performed on an Illumina HiSeq2500 platform (Supplementary Table 1).

### 2.2 Genome assembly and annotation process

#### 2.2.1 Genome size estimation

After trimming, PE Illumina reads of *L. virosa* were used for genome size estimation (∼1,590 million reads; ∼183 Gb). Jellyfish v2.3.0 was used with a k-mer size of 21 to count k-mer frequencies (maximum 1 million count) [20]. The Jellyfish output was used by GenomeScope (v1.0) to estimate haploid genome length, percentage of repetitive DNA, and heterozygosity of the *L. virosa* genome [21].

#### 2.2.2 Assessment of genome completeness

Genome and proteome (annotation) completeness were assessed using BUSCO v5.2.0 with the ‘eudicots_odb10’ dataset [22]. K-mer completeness was assessed with KAT v2.4.1 with a k-mer value of 31 [23].

#### 2.2.3 Genome assembly

PacBio reads were assembled using Canu and then polished by Pilon (v1.20) using Illumina data [24,25]. This assembly was corrected and improved using Bionano optical mapping data. Mis-joins in assembled contigs were corrected using the HiRise pipeline [26]. Since the resulting assembly of Hi-C scaffolding was only 75.2% BUSCO complete, the publicly available—but highly fragmented—assembly for *L. virosa* [18] was used to augment the completeness of our assembly. PE Illumina reads were trimmed before use with Trimmomatic v0.39 ILLUMINACLIP:TruSeq3-PE.fa:2:30:10 LEADING:3 TRAILING:3 SLIDINGWINDOW:4:15 MINLEN:36 [27], and the barcodes of the 10X mate pairs were stripped with Longranger v2.2.2 basic. Before combining the assemblies, we first polished our assembly for a second time with the PE Illumina reads and the 10X mate pair reads (treated as single-end reads) using Pilon v1.24 --changes --diploid --fix all [25]. Mapping of sequencing reads for combining these two assemblies was performed with bwa-mem2 v2.2.1 [28]. Next, we combined our assembly with all sequences >1 kb in the Wei *et al.* [17] assembly. We then aligned all PE Illumina and 10X data to the combined genome. The coverage of this data was used to get the best haplotype representation of the complete genome with purge_haplotigs v1.1.1 (cut-offs were 10, 85, 180) [29]. Since the number of sequences in the resulting assembly increased from 29 to 54,814, we applied several filtering steps to reduce the number of small, uninformative sequences. We filtered out possible mitochondrial and plastid sequences by blasting all sequences to the mitochondrial and plastid NCBI databases (*dd.* 3 August 2021). We filtered out non-*Viridiplantae* sequences as identified by a Blastn search against the NCBI database. Then, we polished the newly added sequences using the same method we used to polish our original genome assembly before (with Pilon v1.24). Based on coverage of PE Illumina and 10X data, we used purge_haplotigs to check whether any duplications were introduced, but since this was not the case, we did not apply purge_haplotigs a second time. For scaffolding the newly added sequences, we mapped the original PacBio data to the genome with minimap2 v2.21 -cxmap-pb [30]. Scaffolding was done with LRScaf v1.12 -misl 3 -t mm [31]. To keep only potential gene coding sequences, we mapped the RNA-seq data with STAR v2.7.7a [32] and removed all sequences lacking a single alignment. Finally, we also removed all sequences smaller than 5 kb.

#### 2.2.4 Repeat annotation

To annotate the repetitive elements in the *L. virosa* genome, a custom library was created by combining different sources: a *de novo* library of TEs created by RepeatModeler (v2.0.1) with -LTRStruct parameter, a *de novo* library of miniature inverted-repeat transposable elements (MITEs) searched by MITE-hunter, and a specific database for the genus *Lactuca* extracted from a combined database of Dfam (20170127) and Repbase (20170127) [33–36]. Then, RepeatMasker (v4.0.7) was used to soft mask the *L. virosa* genome assembly [37]. The same pipeline was also applied to create a TE library and mask the genome assembly of *L. saligna* version 4 (PRJEB35809) and *L. sativa* version 7 (GCF_002870075.2), which were used in reference-based repeatome comparison. The three generated TE libraries were used for a reference-free approach for TE classification (see Individual and comparative clustering analysis of repetitive elements below). The RepeatMasker outputs were further processed to summarize the different categories of repeat elements. Moreover, the LTR elements were extracted from the cross_match output of RepeatMasker and compared to the genome using bedmap.

#### 2.2.5 Gene prediction

Protein encoding genes in the nuclear assembly were annotated using MAKER2, which combines *de novo* gene prediction and homology prediction [38]. rRNA reads were filtered out by SortMeRNA version 4.3.4 [39] using all databases to remove non-coding rRNA. Subsequently, HISAT2 (v2.2.1) was applied to map the remaining reads to the final genome assembly, which includes nuclear sequences, the mitochondrion assembly of CGN013357 (MZ159960.1) and plastid assembly of TKI-404/CGN04683 (CNP0000335 on CNGB) [18,40,41]. The alignment to the nuclear sequences was used as input to BRAKER (version 2) and Stringtie (v2.1.6) to conduct *de novo* gene prediction and transcriptome assembly, respectively, both with default settings [42,43]. The protein alignment was done by BLAST in MAKER2 during the integration with protein databases of *A. thaliana* (Araport11), *L. sativa*, *Helianthus annuus* (HA412.v1), and Uniprot (SwissProt set only: release-2019_10). The predicted transcripts were then filtered using the following criteria: eAED > 0.9 (computed by MAKER2), protein length < 50, identical isoforms, and missing start and stop codon.

#### 2.2.6 Functional annotation

Potential biological function of proteins was inferred using three criteria: i) best-hit matches in SwissProt, TrEMBL using DIAMOND version 2.0.14 at E-value cut-off of 1e-5 [44]; ii) protein domains/structure identified by InterProscan 5.53-87.0 against the Pfam, Coils, Gene3D, PANTHER, SUPERFAMILY, ModiDBLite, and TIGRFAM databases [45,46]; and iii) orthology searches for pathway information were conducted by Kofamscan [47] using a customized HMM database of KEGG orthologs [48] with an E-value cut-off of 1e-5.

### 2.3 Homology analysis

#### 2.3.1 Gene space analysis

To enable a comparison between *L. virosa*, *L. saligna*, and *L. sativa*, we used PanTools v3.4.0 [49] to calculate homologous relationships in a predicted panproteome of these three species. We used the longest isoform for each gene. Based on an optimal distribution of BUSCO genes, we decided to use ‘pantools group -rn 2’ for homology grouping. Subsequent gene classification of the homology groups was also done with PanTools. The number of shared groups were visualized with ComplexUpset [50]. Functional enrichment analyses were performed and visualized for the unique sets of genes with ClusterProfiler v3.18.1 [51].

#### 2.3.2 Synteny detection

MCScanX [52] was utilized to detect syntenic blocks (default settings) among the three *Lactuca* species using the calculated homology groups from PanTools. The interspecific collinearities were visualized using SynVisio [53]. MCScanX was run a second time to detect the tandem arrayed genes using DIAMOND (version 2.0.14) on proteomes for each species.

### 2.4 Individual and comparative clustering analysis of repetitive elements

RepeatExplorer2 on a Galaxy server was used (https://repeatexplorer-elixir.cerit-sc.cz/) to conduct individual and comparative clustering of Illumina PE reads for three *Lactuca* species (*L. sativa*, *L. saligna*, and *L. virosa*) [54]. Re-sequencing data of these three *Lactuca* species were retrieved from ENA database (PRJEB36060). Trimmed FASTQ reads were converted to FASTA format and interlaced prior to the clustering analysis. In addition, a four-letter prefix identity code was added to each sample dataset (i.e., Lsat for *L. sativa*, Lsal for *L. saligna*, and Lvir for *L. virosa*). After a preliminary round, each set of reads was randomly subsampled with a same proportion to maximize the repeat detection and annotation accuracy. For individual analysis, reads representing 20% of the genome size were separately clustered for each *Lactuca* species (i.e., genome proportion = 0.2X, *L. sativa* = 4,166,668 reads, *L. saligna* = 3,833,334 reads, and *L. virosa*= 6,166,668 reads). For comparative analysis, a mixed dataset of reads equal to 0.07X depth for all species was clustered at once (i.e., genome proportion = 0.07X, *L. sativa* = 1,307,006 reads, *L. saligna* = 1,420,966 reads, and *L. virosa*= 2,103,018 reads). For both analyses, the reads were clustered based on the default settings (90% similarity, 55% coverage), and clusters containing more than 0.01% reads were classified at a supercluster level.

After clustering, repeat reads were annotated based on a similarity search to REXdb (protein domain in retrotransposons) using BLAST on a Galaxy server [55]. Additionally, the custom libraries previously created by reference-based searches were utilized to further annotate the repeat clusters (see previous section: Repeat annotation). After annotation, clusters from plastid and mitochondrial origins were identified and excluded for downstream analysis. Next, we quantified different TE categories based on clusters and their connections to superclusters. To characterize the interspecific difference, the clusters resulting from comparative analysis were sorted via hierarchical clustering (ward.D2) using transformed read number [log2(count + 1)] in each cluster for every species.

### 2.5 Analysis of immune gene repertoire

NLRs were searched for in the predicted proteomes of *L. virosa* and *L. sativa,* and retrieved from the *L. saligna* genome [19]. HMMER was used to search Hidden Markov Models (HMMs) profiles obtained from Pfam or the UC Davis database for structural domains of NLR proteins (E-value cut-off = 1e-10): PF00931.23 and NBS_712.hmm (https://niblrrs.ucdavis.edu/At_Rgenes/HMM_Model/HMM_Model_NBS_Ath.html) for the nucleotide-binding (NB) domain; PF01582.20 and PF13676.6 for TIR (TOLL/interleukin-1 receptor); PF05659.11 and PF18052.1 for CC (coiled-coil); and eight HMMs for the LRR (leucine-rich repeat) domain (PF00560.33, PF07723.13, PF07725.13, PF12799.7, PF13306.6, PF13516.6, PF13855.6, PF14580.6). Furthermore, NB and LRR domains identified by InterProScan (see Functional annotation), and CC motifs predicted by Paircoil2 (P scores < 0.025) were combined with the HMMER output [46,56]. The identified NLRs were classified as TNL or CNL based on the presence of either the TIR or CC domain, respectively. To further solve the unclassified NLRs (TNL or CNL), a phylogenetic tree for amino-acid (aa) sequences with NB domains was constructed. First, aa sequences were aligned using HmmerAlign [57]. The alignment was then trimmed by trimAl using -automated1 mode and retained 727 residues for phylogenetic construction [58]. A maximum-likelihood (ML) tree was inferred by IQTREE version 1.6.12 (-m PMB+F+R10) with 1,000 ultrafasta bootstrap (UFBoot) replicates [59]. The phylogenetic tree was visualized and annotated using iTOL v6 [60].

An Inventory of RLKs was also performed for *L. virosa* and *L. sativa*. First HMMER (v3.3.2) was used to search the Pkinase domain (PF00069; E-value cut-off = 1e-10). Then, proteins containing Pkinase were examined for the existence of extracellular domains using HMMER (E-value cut-off = 1e-3) and transmembrane regions using TMHMM (v2.0) and SCAMPI (v2) [61,62].

## 3. Results and Discussion

### 3.1 Genome assembly and annotation

We created a complete and structurally informative genome assembly for *L. virosa* with a total length of 3.45 Gbp (Table 1). Based on a k-mer analysis of Illumina data, we estimated the genome size to be 3.3 Gbp with 73% repeat content and 0.169% heterozygosity rate (Supplementary Figure 1). This predicted size was lower than the previously measured C-value (3.7 Gbp) [16], which might be caused by the large repeat content of *L. virosa* [21]. The long-read assembly was based on PacBio and Illumina data and scaffolded using Bionano and Hi-C data. The longest 12 scaffolds out of the 29 scaffolds comprised 99.8% of the total length (3.3 Gbp) of this first assembly, yet not all chromosomes were reconstructed in full. Therefore, we completed the assembly through additional polishing and leveraging the fragmented, short-read based genome assembly of the same *L. virosa* accession [18] which we combined in a non-redundant way (Supplementary Data 1, Supplementary Figures 2B and 2D). The final combined assembly consisted of 5,855 contigs spanning a total of 3.45 Gbp with an N90 (L90) score of 116,478,781 (10) (Supplementary Figure 2C; Supplementary Table 2). The BUSCO completeness score was 96.2% (the duplication score was 4.5%; Supplementary Table 3; Supplementary Figure 2D).

**Table 1.**
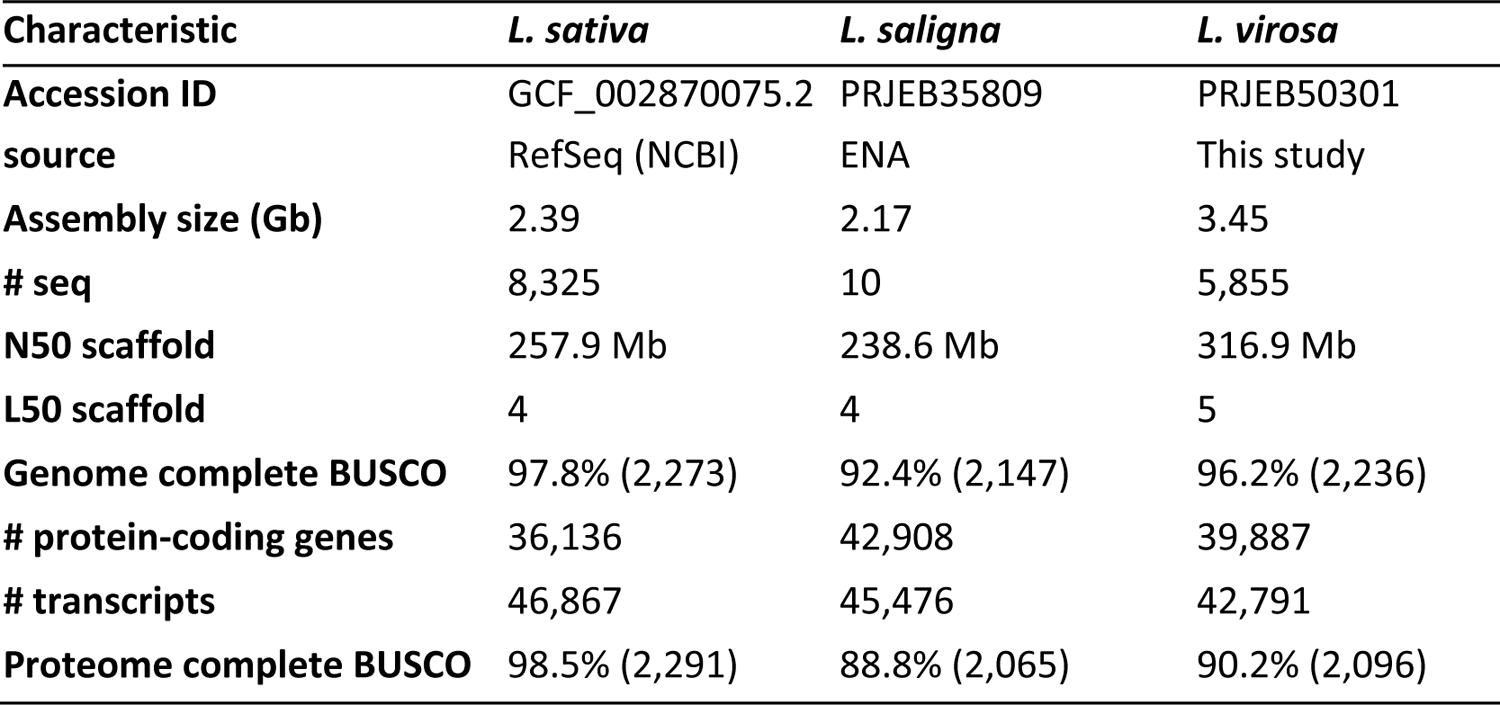
Summary of assemblies for *Lactuca* spp. in this paper.

| Characteristic | *L. sativa* | *L. saligna* | *L. virosa* |
| --- | --- | --- | --- |
| Accession ID | GCF_002870075.2 | PRJEB35809 | PRJEB50301 |
| source | RefSeq (NCBI) | ENA | This study |
| Assembly size (Gb) | 2.39 | 2.17 | 3.45 |
| # seq | 8,325 | 10 | 5,855 |
| N50 scaffold | 257.9 Mb | 238.6 Mb | 316.9 Mb |
| L50 scaffold | 4 | 4 | 5 |
| Genome complete BUSCO | 97.8% (2,273) | 92.4% (2,147) | 96.2% (2,236) |
| # protein-coding genes | 36,136 | 42,908 | 39,887 |
| # transcripts | 46,867 | 45,476 | 42,791 |
| Proteome complete BUSCO | 98.5% (2,291) | 88.8% (2,065) | 90.2% (2,096) |

Based on both expression and orthology evidence, 39,887 protein-coding genes with a total of 42,791 transcripts were annotated. We mapped RNA-seq data from root, leaf, and flower tissue to the genome assembly to support *de novo* gene prediction. Next, we aligned protein sequences of model plant species to the genome and used MAKER for merging all gene predictions. We filtered the predicted genes to only retain annotations that were in accordance with the provided evidence. The BUSCO score on the resulting proteome was 90.2%, indicating a high level of completeness. Furthermore, we were able to predict functional domains in 93% (37,106) of the genes for various databases (Supplementary Table 4; Supplementary Data 2A - B). This structural and functional annotation is vital for the biological interpretation of *L. virosa* data.

### 3.2 Homology grouping of three representative *Lactuca* spp

Even though the genome size of *L. virosa* is substantially larger than *L. sativa* and *L. saligna*, the number of genes annotated across species was similar (Table 1). A comparison of *L. virosa* with *L. saligna* and *L. sativa* showed that about half of the homology groups are shared across *Lactuca* (Figure 1; Supplementary Data 2C). These 17,741 homology groups in *Lactuca* contained 19,270 *L. virosa* genes, meaning that about half of the *L. virosa* genes are part of the core *Lactuca* genome. This is comparable to what was found in other interspecies comparisons. For example, in rice ∼62% of core genes were reported between two species [63], and in *Raphanus*, ∼50% of core genes were reported among 11 accessions belonging to two species [64]. Both *L. virosa* and *L. saligna* share fewer homologous genes with each other than with *L. sativa*. This stresses the importance of wild species in breeding as they contain a large pool of novel genes. The large, unique genomes of both *L. virosa* and *L. saligna* indicate that these wild species are rich sources of genetic diversity that thus far has been unexploited for lettuce breeding. We performed a functional annotation for the proteomes of the three species with InterProScan to perform functional enrichment for the unique content of *L. virosa* (15,048 genes; Supplementary Data 2D). The InterProScan domain enrichment found disease resistance proteins to be among the set of significantly enriched domains (Supplementary Figure 3). Therefore, the genome of *L. virosa* is a resource for potential novel genes needed for resilience breeding in lettuce. Furthermore, it will be relevant to sequence and produce high-quality assemblies of other wild relatives of lettuce, such as *L. georgica*, *L. serriola*, and *L. aculeata,* to obtain an overview of the entire *Lactuca* gene space [18,65]. Using high-quality genetic resources will enable the construction of a comprehensive pangenome that covers the variation in the genus *Lactuca*.

**Figure 1.**
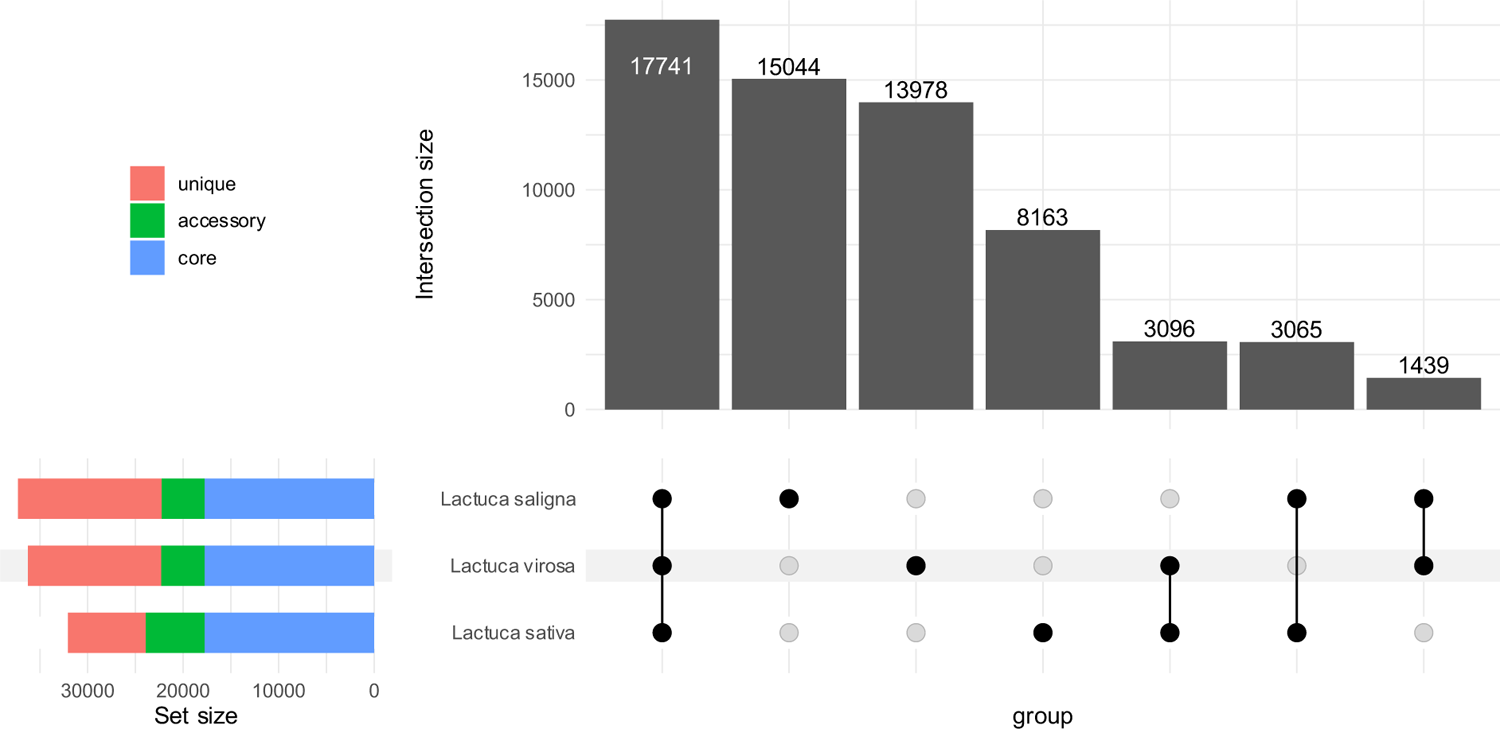
Overview of homology grouping for *L. sativa*, *L. saligna*, and *L. virosa* in an upset plot. The numbers on bars are groups of homologous genes. In total, there are 62,526 homology groups.

### 3.3 Synteny detection between three *Lactuca* spp. via comparative genomics

By synteny detection of homologous pairs, we identified major chromosomal inversions between the three *Lactuca* genomes. Overall, there was whole-genome collinearity (synteny) among *Lactuca* species (Figure 2D). Based on the collinearity, we determined the major 12 scaffolds that comprised 96% (3.30 Gbp) of the total genome assembly (Supplementary Table 5). Compared to the *L. sativa* genome, three species-specific inversions were identified on different chromosomes (Chr; Figure 3). Two of the three inversions that were previously described between *L. saligna* and *L. sativa* were validated and further characterized: one is specific to *L. saligna* on Chr5 and one is specific to *L. sativa* on Chr8 [19]. Furthermore, synteny also revealed a large inversion specific to *L. virosa* on tentative Chr7 (Scaffold8) (Figure 3; Supplementary Table 5). These inversions could hamper genetic mapping of interesting traits and further introgression. The syntenic patterns between *L. virosa* Chr9 (scaffold7) and the other two species showed complicated inverted and translocated regions, which might be due to a reversed-joining from Hi-C scaffolding (Supplementary Figure 4).

**Figure 2.**
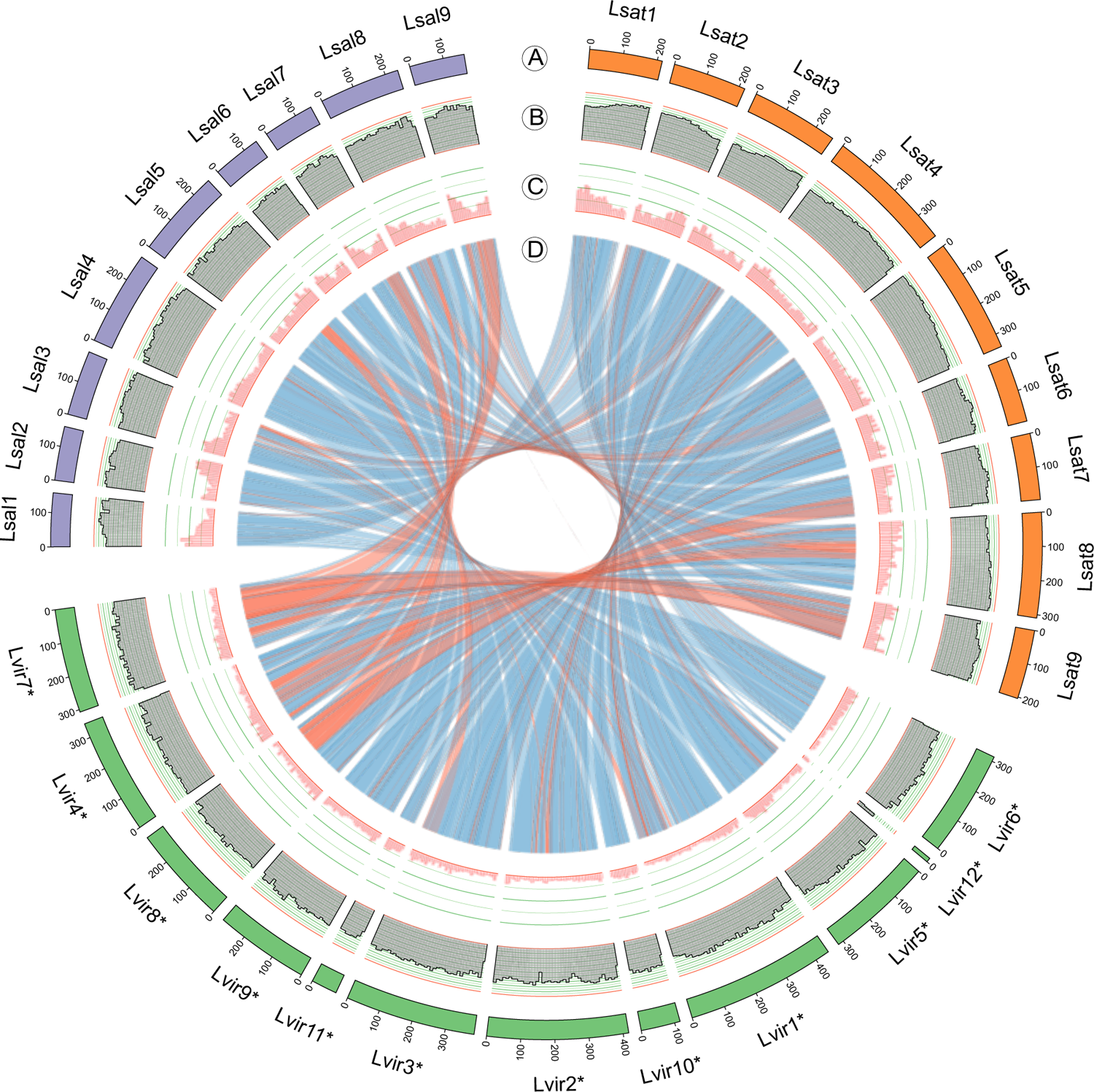
Circos plot of *L. virosa* genome compared with the *L. sativa* and *L. saligna* genomes. **A,** For each of the three genomes (Lsat: *L. sativa*; Lvir: *L. virosa*; Lsal: *L. saligna*), only sequences larger than 1 Mbp are shown. For *L. sativa* and *L. saligna*, the number of sequences correspond to their chromosomes. Since *L. virosa* is near-chromosome level, its sequence numbers were indicated with an asterisk (*). The sequences were sorted based on its collinearity to other *Lactuca* species, and some of the sequences for *L. virosa* are inverted to match the orientation in previously published genomes (**D**); the sequence coordinates (in Mbp) show this. **B,** The repeat density for each sequence is calculated per 10 Mbp and shown here as a fraction. Since the genome assembly for *L. virosa* has more N bases, repeats are more difficult to find than in the other two genomes. The scale ranges between 0 and 1. **C,** The gene density for each sequence is calculated per 10 Mbp and shown here as a fraction. As the three genomes contain approximately the same number of genes but their genome sizes differ, *L. virosa* has a lower overall gene density. The scale goes from 0 to 0.2. **D,** Synteny between the three genomes. Inversions are shown in red as opposed to non-inverted syntenic blocks, which are shown in blue.

**Figure 3.**
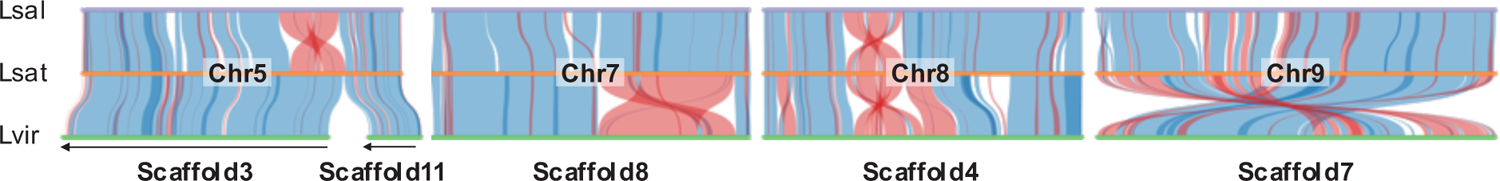
Synteny discloses species-specific inversions across three *Lactuca* species. Through genomic comparison, major interspecific inversions (red) were identified among the reference genomes of three *Lactuca* species. Here, the synteny in four sets of scaffolds/chromosomes reveal species-specific inversions: *L. saligna* (Lsal: purple), *L. sativa* (Lsat: orange), and *L. virosa* (Lvir: green). The chromosome numbers are labelled in the middle. Black arrows at the bottom indicate reversed scaffolds in the *L. virosa* assembly. Supported by Supplementary Table 5.

### 3.4 Comparative repeatomics between three *Lactuca* spp. via reference-based and reference-free approaches

In the three reference assemblies, we annotated repeat elements and classified them into TEs and other repeats (Supplementary Data 3A). The genomes of all three *Lactuca* species contained a major proportion of TEs, in agreement with previous studies (Supplementary Table 6) [13]. Intriguingly, the TE content of the *L. virosa* (60%) assembly is lower than that of both *L. sativa* (74%) and *L. saligna* (77%), whereas a higher TE content was expected in *L. virosa*. After excluding the N content of each genome, the percentage of TEs for all *Lactuca* genomes exceeded 80% (Supplementary Figure 5; Supplementary Table 6). Moreover, LTR TEs represented more than 50% of the three *Lactuca* genomes (excluding N content). We further characterized LTRs in the *L. virosa* genome by determining their genomic position. Almost all identified LTRs (99%) were located in the intergenic regions and their density gradually decreased nearing a genic region (Supplementary Figure 6). Moreover, the non-repeat regions (i.e., unmasked length, excluding N content) were similar in all species, regardless of the change in repeat length (Supplementary Table 6). To conclude, this reference-based repeat annotation showed that TEs are the most abundant components of *Lactuca* spp. genomes. However, genome incompleteness and N content of the reference genome assemblies hamper a precise estimation of TEs.

In addition to the reference-based repeat annotation, we also classified repeat components and estimated their composition for the three *Lactuca* spp. in a reference-free way by annotating the clusters of repetitive re-sequencing short reads (Supplementary Table 7). First, we performed an individual analysis for three *Lactuca* species with a read depth of 0.2x. In total, we found that *L. virosa* had the highest percentage of repeated reads assembled as clusters (82%). For top clusters (cluster size > 0.01% analyzed reads), the percentage of repeated reads was in line with the estimated genome size for three *Lactuca* species: *L. virosa* (74%), *L. sativa* (68%), and *L. saligna* (65%). After curation, we calculated the genomic proportion of different types of TEs based on the annotated read clusters (Supplementary Table 8). In all species, more than 60% of repeated sequences were annotated as TEs, which comprised almost 100% of Class I LTRs. Among them, *L. virosa* had the highest amount of LTRs (68.34%) followed by *L. sativa* (61.24%) and *L. saligna* (58.64%). It is likely that the overall genome size expansion of *L. virosa* was driven by TEs.

To explore this further, we identified the major differences in TEs represented by read clusters between three *Lactuca* species via another comparative analysis (RepeatExplorer). This approach used mixed reads at the same depth (0.07x) for every species (Supplementary Table 7). Compared to the individual mode, the clusters resulting from the comparative analysis contained the read number from each species for each cluster, i.e., a cluster matrix for three *Lactuca* species (Supplementary Data 3B). For example, cluster 10 was annotated as LTR/Gypsy and mainly composed of *L. virosa* reads (Supplementary Figure 7). After curation, the total repeat content of the top clusters was 69.45% for mixed data, which was higher than *L. sativa* (63.54%) and *L. saligna* (61.75%) and lower than *L. virosa* from individual analysis (70.40%), indicating that *L. virosa* carries a higher percentage of repeats than the other two *Lactuca* species (Supplementary Table 8).

Based on the comparative analysis, an in-depth cluster analysis revealed that the LTR proliferation in *L. virosa* drove its genome expansion (Supplementary Data 3C and 3D). The heatmap of hierarchical clustering shows six groups that were either dominated (D) by one of the three *Lactuca* species: *L. sativa* (Lsat), *L. saligna* (Lsal), or *L. virosa* (Lvir) (Figure 4A: left). The bar plot in Figure 4A (right) further decomposes the read sources for each group. The Lvir_D2 group, dominated by *L. virosa* reads, was also the largest group in the hierarchical clustering analysis. Further investigation of the genomic proportion for clusters in the Lvir_D2 group was carried out. Collectively, the clusters in Lvir_D2 comprised 50.05% of the reads and were significantly larger than the other five groups. This Lvir_D2 group is dominated by *L. virosa*, contributing approximately two times as many reads as *L. sativa* and *L. saligna*. Furthermore, nearly all read clusters in Lvir_D2 were annotated as LTR (48.47% of analyzed reads) and mainly represented by two LTR sub-families: Gypsy (27.31%) and Copia (20.46%) (Supplementary Table 9). Additionally, the subgroups Tekay and Angela were the primary elements for the Gypsy and Copia clusters within the Lvir_D2 group (Figure 4B; Supplementary Table 9).

**Figure 4.**
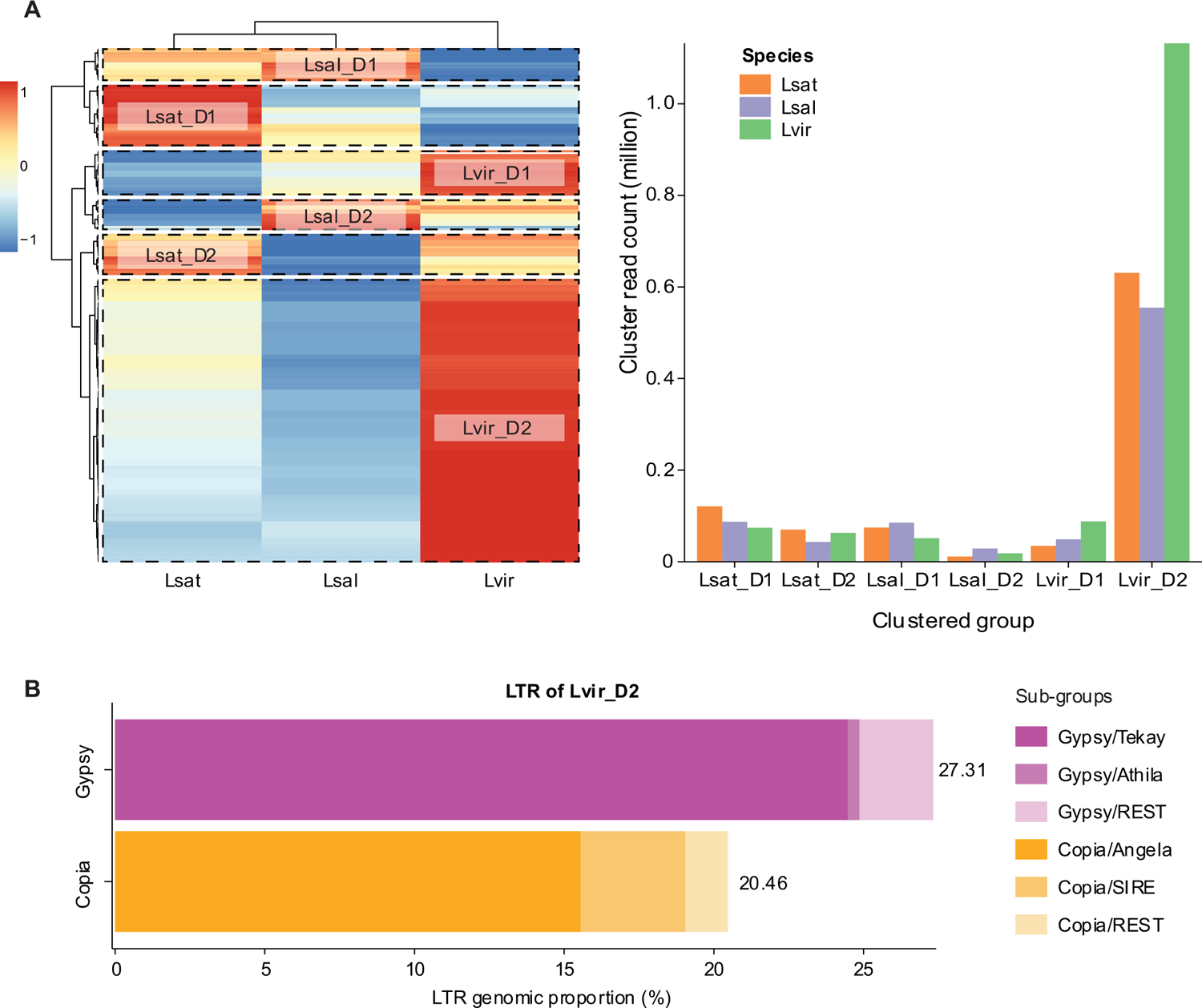
Proliferation of long-terminal repeats (LTR) drives the expansion of the *L. virosa* genome. Read clusters assembled by RepeatExplorer2 using a mix of resequencing data (coverage = 0.07x) from three *Lactuca* species references to detect the major difference of repeat elements. **A,** Heatmap shows the scaled read-count of individual cluster (row) for each species (column). Clusters and species were sorted by hierarchical clustering. Six groups (squared by dash lines) were dominated by reads either from *L. sativa* (Lsat), *L. saligna* (Lsal), or *L. virosa* (Lvir) and suffixed with a D (dominant). Bar plot shows the size (y-axis) of six clustered groups (x-axis) for each species: *L. sativa* (orange), *L. saligna* (purple), and *L. virosa* (green). **B,** Stacked bar chart shows the composition of sub-groups for the two major LTR superfamilies: Gypsy (gradient purple) and Copia (gradient yellow). Supported by Supplementary Table 9.

*L. virosa* is estimated to have a significantly larger genome (3.7 Gbp) than *L. sativa* (2.5 Gbp) and *L. saligna* (2.3 Gbp) [16]. TEs have been shown to drive plant genome expansion [17]; for example, within the genus of rice [66–68]. Based on our combined findings, we conclude that the subgroups of transposon LTR, Tekay in Gypsy, and Angela in Copia drove the genome expansion of *L. virosa*.

### 3.5 Comparison of NLR and RLK genes between three *Lactuca* spp

Besides the difference within TEs, there is also sizable variation in the number of genes as shown by the homology grouping (accessory/unique genes) among these three *Lactuca* species (Supplementary Figure 3), which might convey resilience to important traits like resistance against various pathogens or pests. In our previous study, an extensive search of resistance genes was performed for lettuce and its wild relative *L. saligna* [19]. Using the new *L. virosa* assembly, we identified and classified immunity-related genes encoding NLR and RLK proteins for *L. virosa* and compared them to *L. sativa* and *L*. *saligna*.

The *L. sativa* genome was found to have the highest number of NLRs (385), followed by *L. saligna* (323) and *L. virosa* (309) (Table 2; Supplementary Table 10). In association with the homology grouping, a Venn diagram showed that the NLRs identified in three *Lactuca* spp. are highly diverged, where more than 50% of NLRs in each species belong to specific homology groups (Figure 5A: left; Supplementary Data 5A). This observation is in line with our enrichment study of homologs specific to *L. virosa*, where InterProScan domains were significantly enriched with terms related to NLR proteins (Supplementary Figure 3). Furthermore, NLR proteins were classified into TNL and CNL types based on the N-terminal domain (TIR or CC domain, respectively) and curated by the phylogeny of a nucleotide-binding (NB) domain alignment (Supplementary Figure 8; Supplementary Data 4A - B). The difference between *L. sativa*, *L. saligna*, and *L. virosa* was mainly contributed by *TNL* genes (227 vs 184 and 180), and the difference between *L. saligna* and *L. virosa* can be explained by the *CNL* type (139 vs 162). Due to the unequal completeness of the proteomes, we applied the ratio of complete BUSCOs for proteomes as a benchmark to anticipate whether *NLR genes* expand or contract between the three *Lactuca* spp.: *L. sativa* (2,291), *L. saligna* (2,065), and *L. virosa* (2,096). The ratio of BUSCOs (1.10 : 1.00 : 1.02) reflects the NLR ratio across species (1.25 : 1.05 : 1.00), where *L. sativa* showed a slight inflation. For different NLR types, the number of CNLs was similar in the examined species *sativa*, *L. saligna*, and *L. virosa* (1.14 : 1.00 : 1.06); however, the ratio of TNL numbers highly deviated from the BUSCO ratio (1.41 : 1.14 : 1.00; Supplementary Table 10). Such comparison suggests an expansion of *NLR*s in *L. sativa*, which is possibly caused by tandem duplication events as in most studied angiosperms [69]. This hypothesis is supported by whole-genome search of tandem duplicates (TDs) clusters between three *Lactuca* spp. genomes (Supplementary Data 5B). The number of TDs encoding NLRs in *L. sativa* (121) was approximately two-times larger than that in *L. saligna* (61) and *L. virosa* (76), which principally explains the number difference among the three species (Figure 5B: left). In addition to tandem duplication, transposon activities (e.g. LTRs) could also greatly elevate the number of *NLR*s by retroduplication as reported in the chili genome [70]. The retroduplicated *NLR*s could partially explain the lineage-specific homologs among *Lactuca* species (Figure 5A: left).

**Figure 5.**
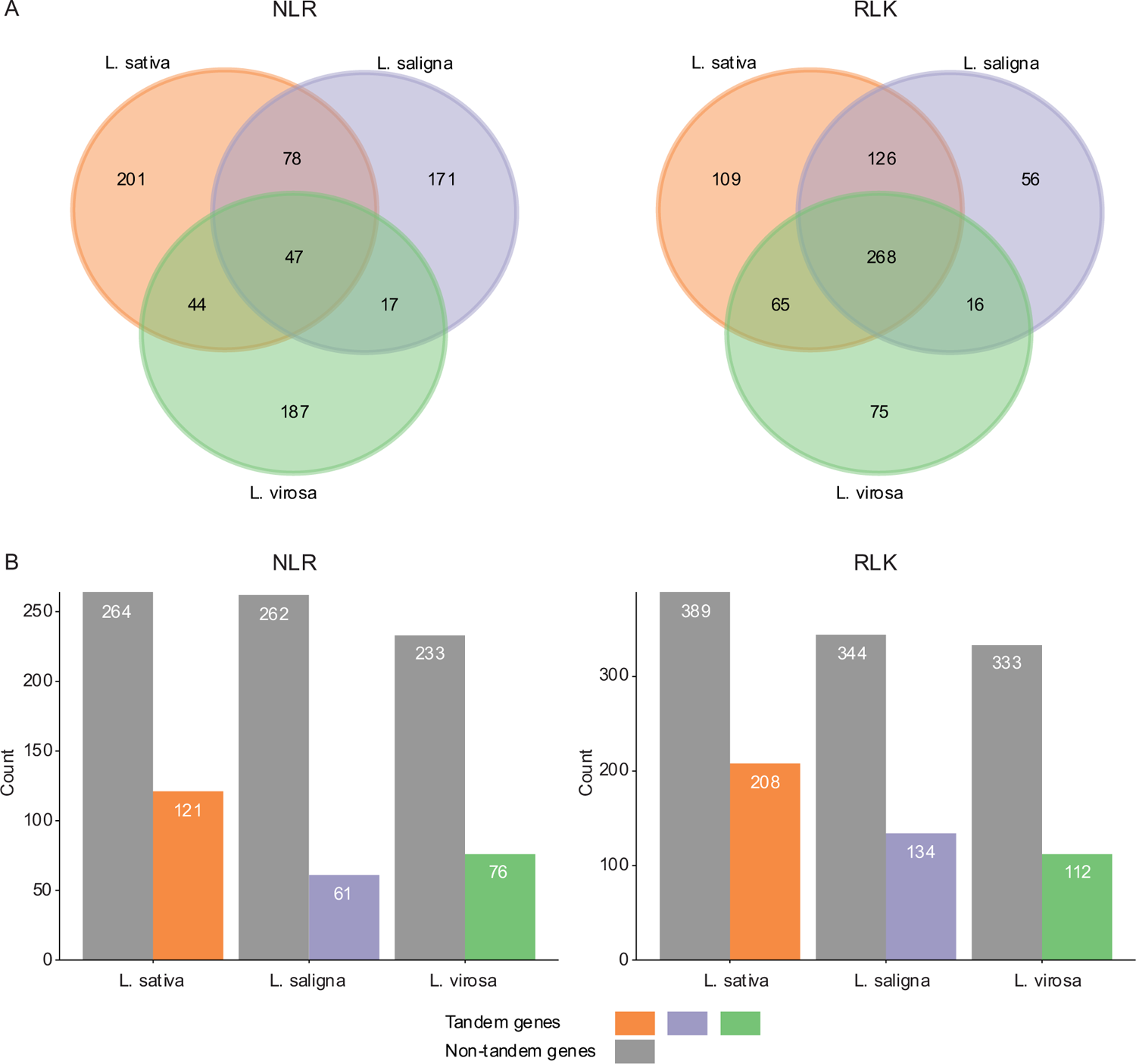
Homology and tandem duplication relationships of immune genes across *Lactuca* species. **A,** Venn diagrams of homology groups for nucleotide-binding leucine-rich repeats (*NLRs*) and receptor-like kinases (*RLK*s) in *Lactuca* spp. Homology grouping was done by PanTools. **B,** Bar plots show the count of tandem (colors) and non-tandem (gray) *NLRs* and *RLKs* in three *Lactuca* species. Tandem arrayed genes were identified by MCScanX. Supported by Supplementary Data 5.

**Table 2.** Identification and classification of candidate immunity related genes for *Lactuca spp*.

| Immune genes |  | Species |  |  |
| --- | --- | --- | --- | --- |
| Family | Classification | *L. sativa* | *L. saligna* | *L. virosa* |
| *NLR* | CNL* | 158 | 139 | 148 |
|  | TNL | 227 | 184 | 161 |
|  | Total | 385 | 323 | 309 |
| *RLK\*\** | Rcc1-RK | 5 | 5 | 2 |
|  | WAK | 61 | 48 | 36 |
|  | G-LecRK | 132 | 79 | 70 |
|  | L-LecRK | 31 | 29 | 21 |
|  | C-LecRK | 1 | 1 | 1 |
|  | CRK | 41 | 35 | 38 |
|  | Malectin-RK | 55 | 55 | 32 |
|  | LysM-RK | 12 | 12 | 11 |
|  | LRR-RK | 258 | 213 | 233 |
|  | PERK | 1 | 1 | 1 |
|  | Total | 597 | 478 | 445 |

\* RPW8 and Rx\_N type of CNL included in this study.
\*\* RLK classification based on extracellular domain (Supplementary Table 12 and 13).

We next identified RLK proteins by searching for the extracellular, transmembrane, and intracellular domains. Then, resulting RLKs were classified into nine types based on their extracellular and kinase domains (Supplementary Table 11-12; Supplementary Data 4C-D). Like NLRs, we found more genes encoding RLK proteins in the *L. sativa* (597) genome assembly than in *L. saligna* (478) or *L. virosa* (445; Table 2). Sequence similarity shows that *RLK*s were much more conserved in *Lactuca* spp. compared to NLRs, where 70% of *RLK*s in each *Lactuca* species were homologous to another RLK from at least one sister species (Figure 5A: right). Compared to the BUSCO completeness, the RLK ratio (1.25 : 1.00 : 1.00) showed an increase of *RLKs* in *L. sativa*, suggesting a possible expansion of the *RLK* family. The expansion in *L. sativa* was majorly contributed by *G-LecRK*, followed by *Malectin-RK* and *WAK*, while other types of *RLK*s were either similar in all species or slightly inflated in *L. sativa*. The extra *G-LecRK* and *WAK* copies might confer specific immunity in *L. sativa*. For example, G-LecRK and WAK can both mediate resistance to *Phytophthora spp.* (oomycete) in tobacco and melon plants [71,72]. On the contrary, the expansion of *Malectin-RK* might benefit pathogen invasion in *L. sativa*, like the increased susceptibility to *Hyaloperonospora arabidopsidis* (oomycete) observed in *Arabidopsis* [73]. Similar to *NLR*s, *RLK*s also commonly expand via tandem duplications. For example, a *G-LecRK* expansion was reported in soybean [74,75]. The number of tandem arrayed *RLK*s in *L. sativa* was 1.5 and 1.9 times that of the *RLK*s in *L. saligna* and *L. virosa*, respectively, which constitutes more than 60% of the difference between *L. sativa* and other two species (Figure 5B: right; Supplementary Data 5B). Especially for *G-LecRK*, the number of tandem genes appeared more than doubled in *L. sativa* (Supplementary Data 5B).

## 4. Conclusions

Here, we present a near chromosome-level genome assembly for *L. virosa* (accession CGN04683) that has a high completeness. As a representative of the tertiary lettuce gene pool, this *L. virosa* genome assembly enables comparisons with *L. sativa* of the primary gene pool and *L. saligna* of the secondary gene pool. For gene content, *L. virosa* harbors a large number of genes absent from *L. saligna* and *L. sativa* and may thus constitute an important source of novel genes for lettuce breeding. Based on synteny, a three-way genome comparison uncovered species-specific major inversions. These inversions should be considered as likely barriers to gene introgression in future breeding. In addition, we demonstrated the genome expansion in *L. virosa* is driven by the proliferation of LTR elements. An assembly-based comparison of *NLR* and *RLK* genes between *Lactuca* spp. found more immune system-related genes in the *L. sativa* genome than in those of the *L. virosa* and *L. saligna* genomes. These findings may contribute to future research on gene expression and regulation in *L. virosa.* Using this novel genome assembly, researchers can subsequently study the genetic variation in *L. virosa* populations to fully release its potential for lettuce breeding.

### Data availability

The genome assembly of *L. virosa*, is available under the BioProject PRJEB50301 (and available under CAKMRJ010000000.1 from ENA). All raw sequencing reads have been deposited in the ENA database under BioProject PRJEB56289. This includes the Illumina, PacBio, Bionano, and Hi-C whole-genome sequences as well as RNA sequencing data for genome annotation.

### CRediT authorship contribution statement

**Wei Xiong**: Software, Formal analysis, Investigation, Data Curation, Writing - Original Draft, Writing - Review & Editing, Visualization. **Dirk-Jan M. van Workum**: Software, Formal analysis, Data Curation, Writing - Original Draft, Writing - Review & Editing, Visualization. **Lidija Berke**: Methodology, Software, Investigation, Formal analysis, Writing - Review & Editing. **Linda V. Bakker**: Software, Formal analysis. **Elio Schijlen**: Resources. **Frank F.M. Becker**: Methodology. **Henri van de Geest**: Software, Formal analysis. **Sander Peters**: Conceptualization, Methodology, Writing - Review & Editing. **Richard Michelmore**: Resources, Writing - Review & Editing. **Rob van Treuren**: Conceptualization, Resources, Writing - Review & Editing. **Marieke Jeuken**: Conceptualization, Methodology, Investigation, Resources, Writing - Review & Editing. **Sandra Smit**: Software, Supervision, Writing - Review & Editing. **M. Eric Schranz**: Conceptualization, Methodology, Supervision, Writing - Review & Editing, Project administration, Funding acquisition.

## Acknowledgement

We thank Jiri Macas and Floris Breman for their help with RepeatExplorer2 analysis. We also thank Elizabeth Marie Georgian (UC Davis) for writing support.

## Supplementary Figures

**Supplementary Figure 1.**
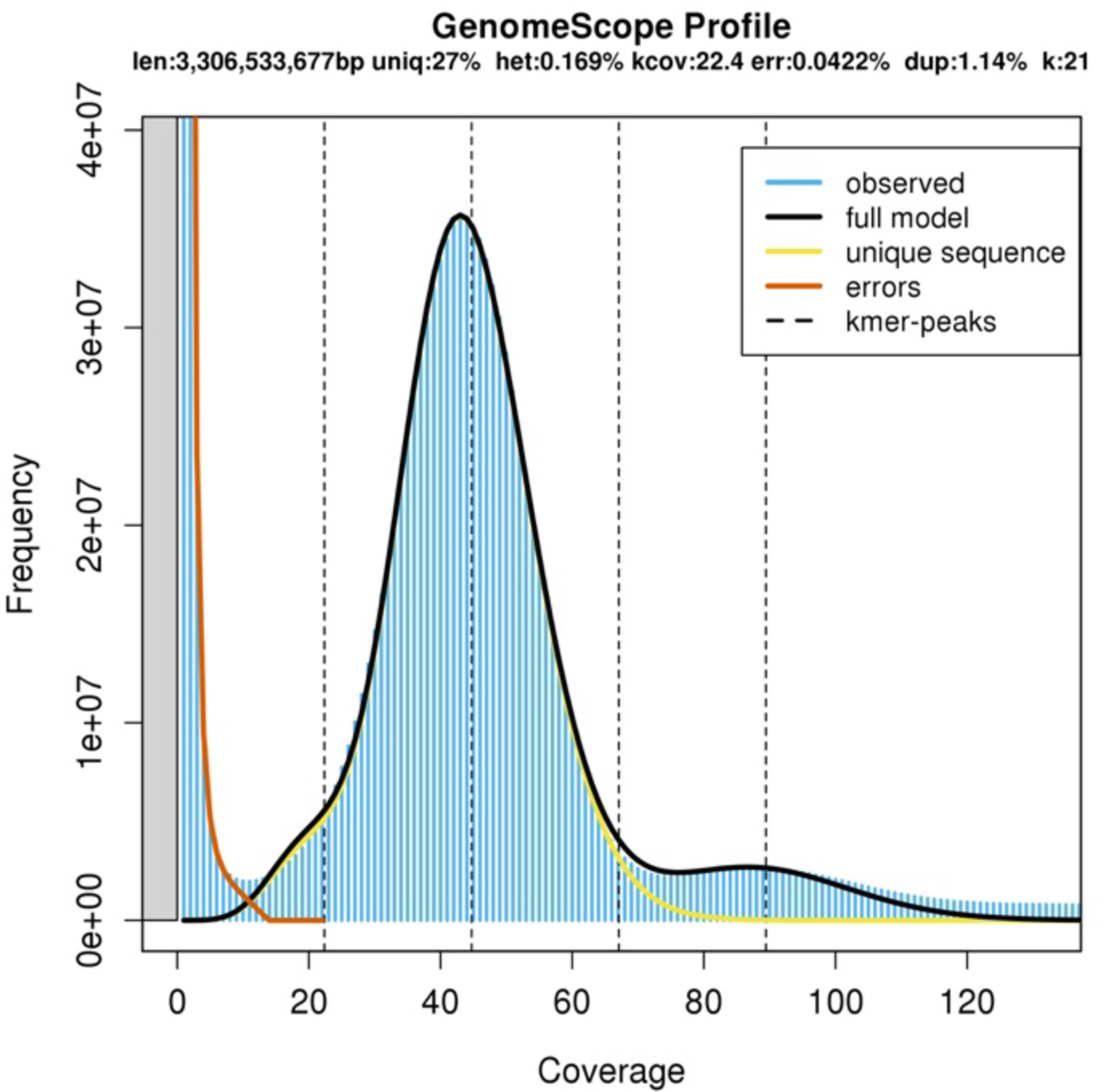
Genome size estimation of *L. virosa* by GenomeScope. The 21-mers were counted by Jellyfish, and output was taken by GenomeScope to estimate the genome size of *L. virosa*. The frequency (y-axis) and sequencing depth (x-axis) of 21-mers are plotted. The genome size (3.3 Gbp) was estimated by the highest peak depth. len: Genome haploid length; uniq: genome unique length; het: heterozygosity rate; kcov: k-mer coverage; err: read error rate; dup: average rate of read duplication; k: k-mer length.

**Supplementary Figure 2.**
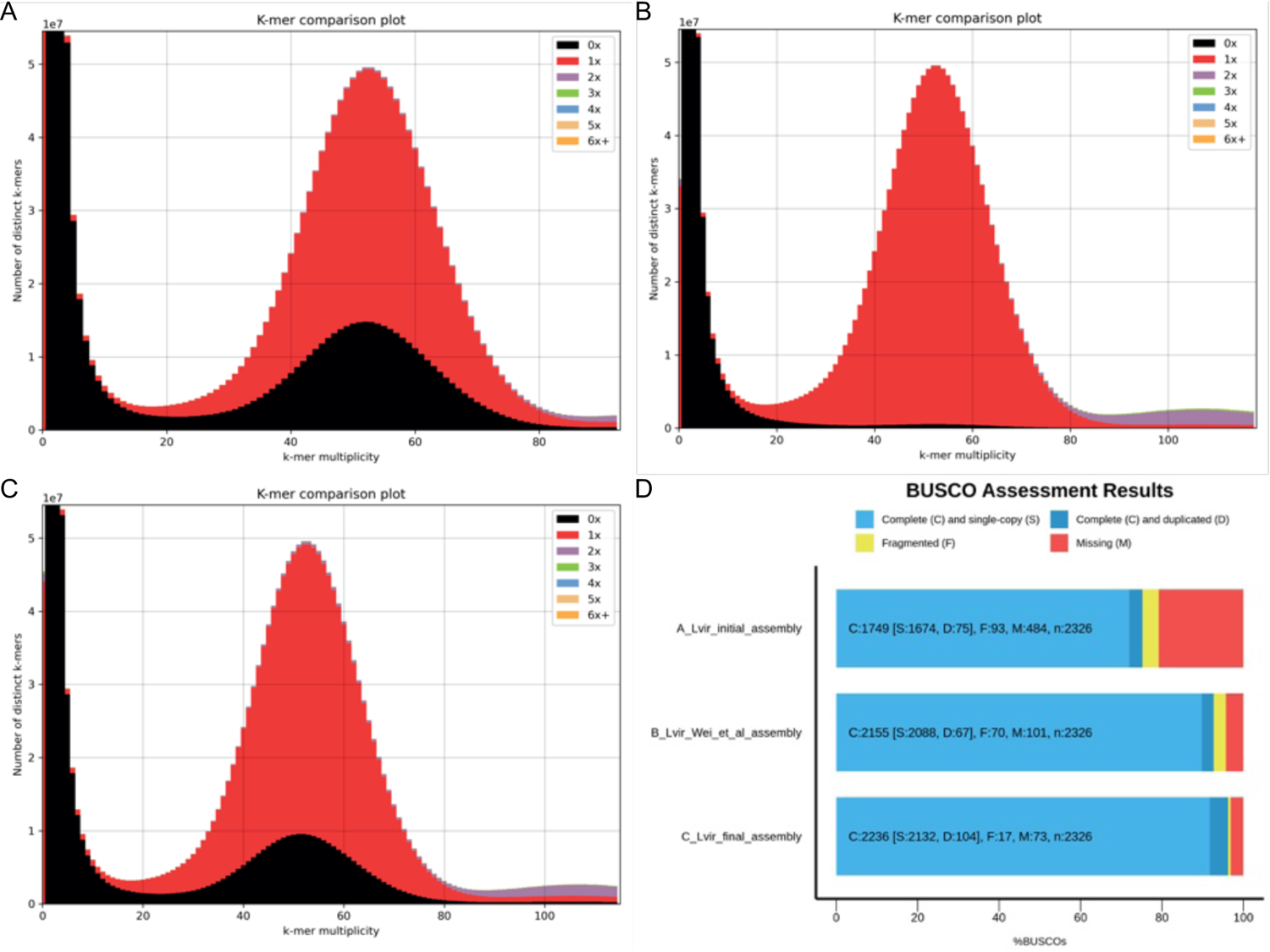
K-mer and BUSCO completeness plots for the first *L. virosa* assembly. **A**, The initial assembly. **B**, The BGI *L. virosa* assembly. **C**, The final *L. virosa* assembly. **D**, BUSCO analysis of these three genomes. For the comparisons in **A-C**, all k-mers in the Illumina data were counted and subsequently colored according to their occurrence in the genome assembly. For **D**, eudicots_odb10 was used.

**Supplementary Figure 3.**
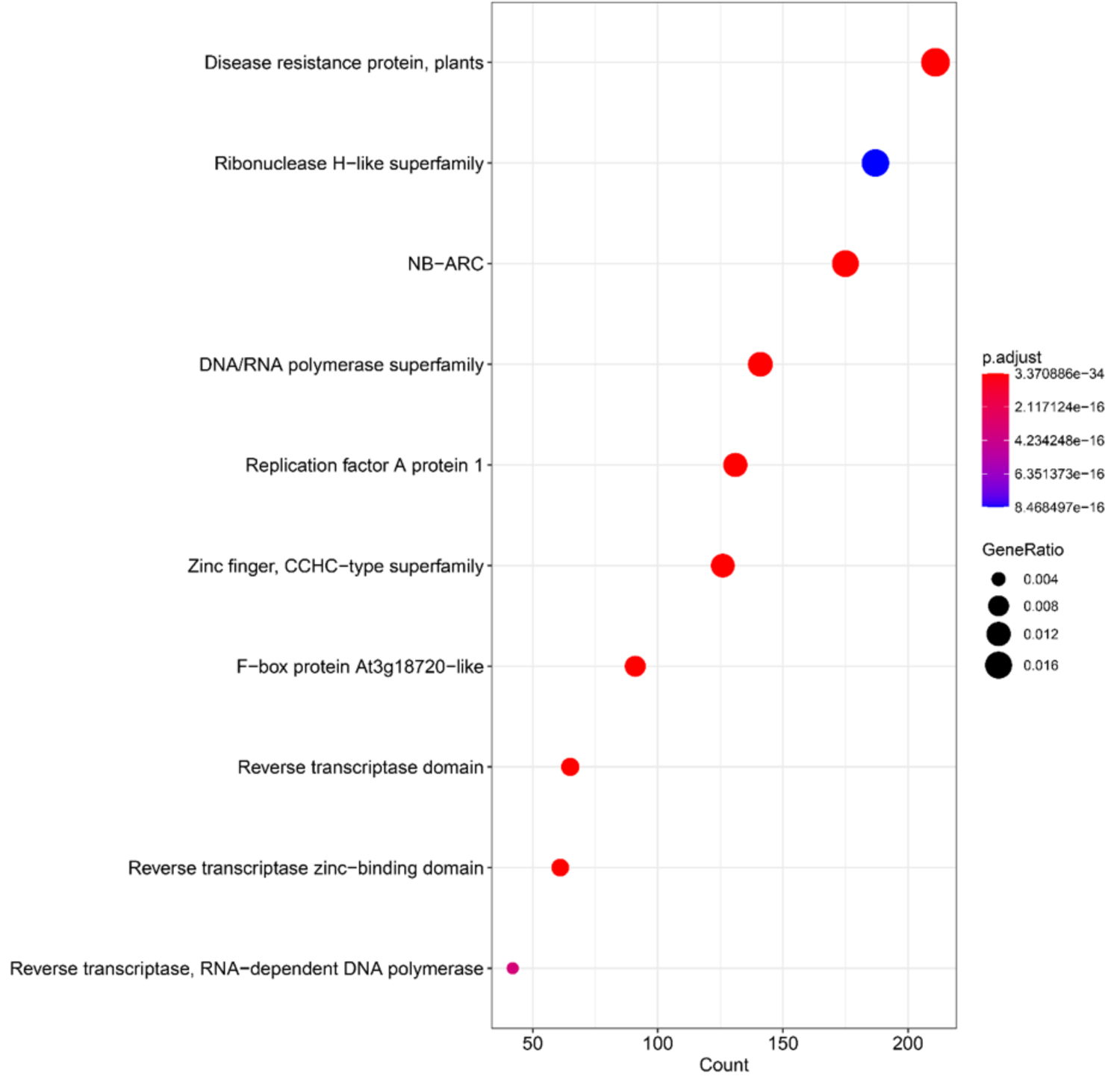
Functional enrichment of genes unique to *L. virosa*. The top 10 most significant hits from an InterPro domain enrichment of all unique *L. virosa* genes according to homology grouping. For the full results, see Supplementary Data 2D.

**Supplementary Figure 4.**
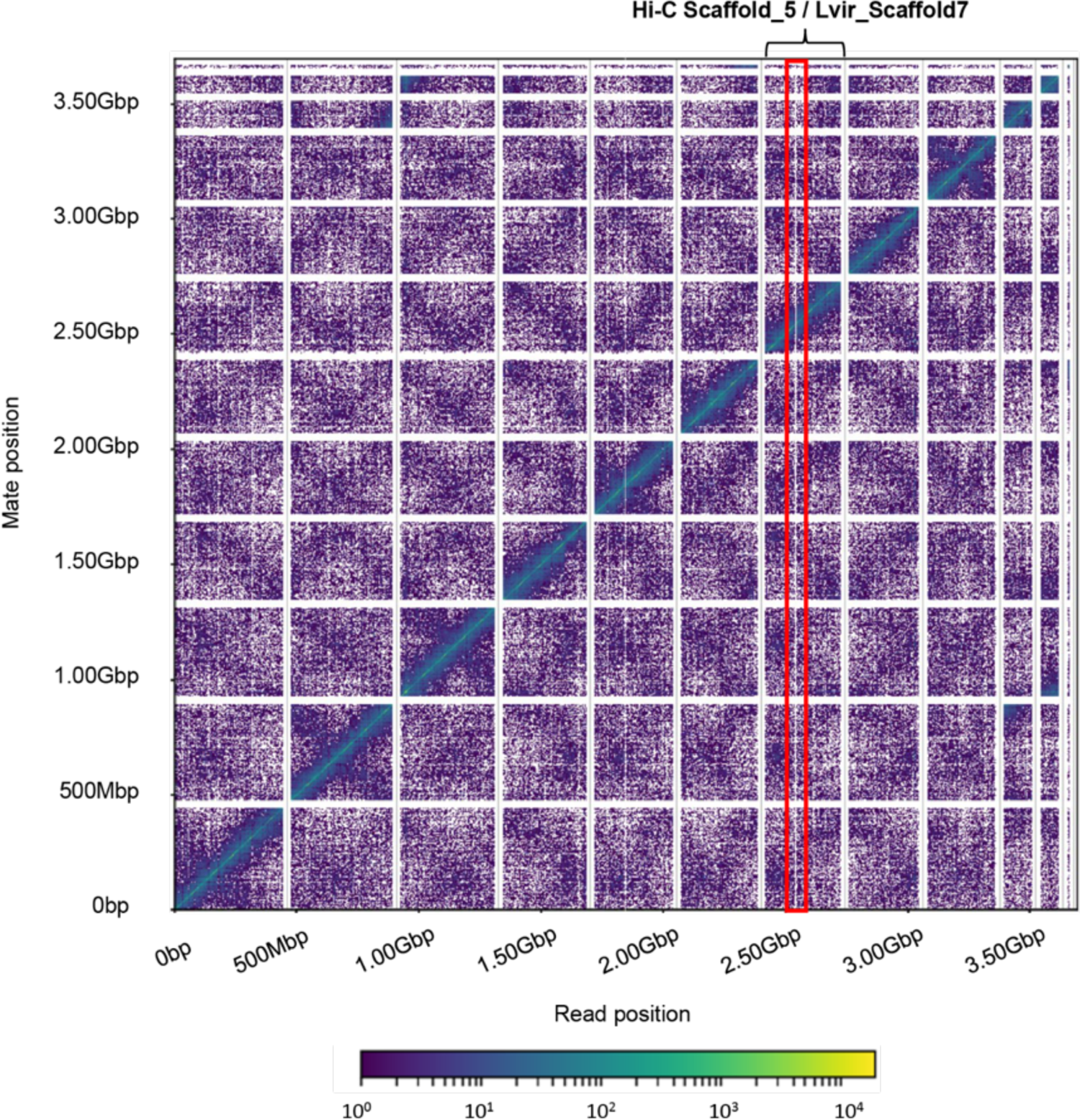
Link density histogram (generated by Dovetail Genomics). In this figure, the x and y axes give the mapping positions of the first and second read in the read pair, respectively, grouped into bins. The color of each square gives the number of read pairs within that bin. White vertical and black horizontal lines have been added to show the borders between scaffolds. Scaffolds less than 1 Mb are excluded. The red rectangle shows a potential breakage in the middle of Scaffold 5 (Hi-C assembly)/Lvir_scaffold7 (final assembly) for *L. virosa*.

**Supplementary Figure 5.**
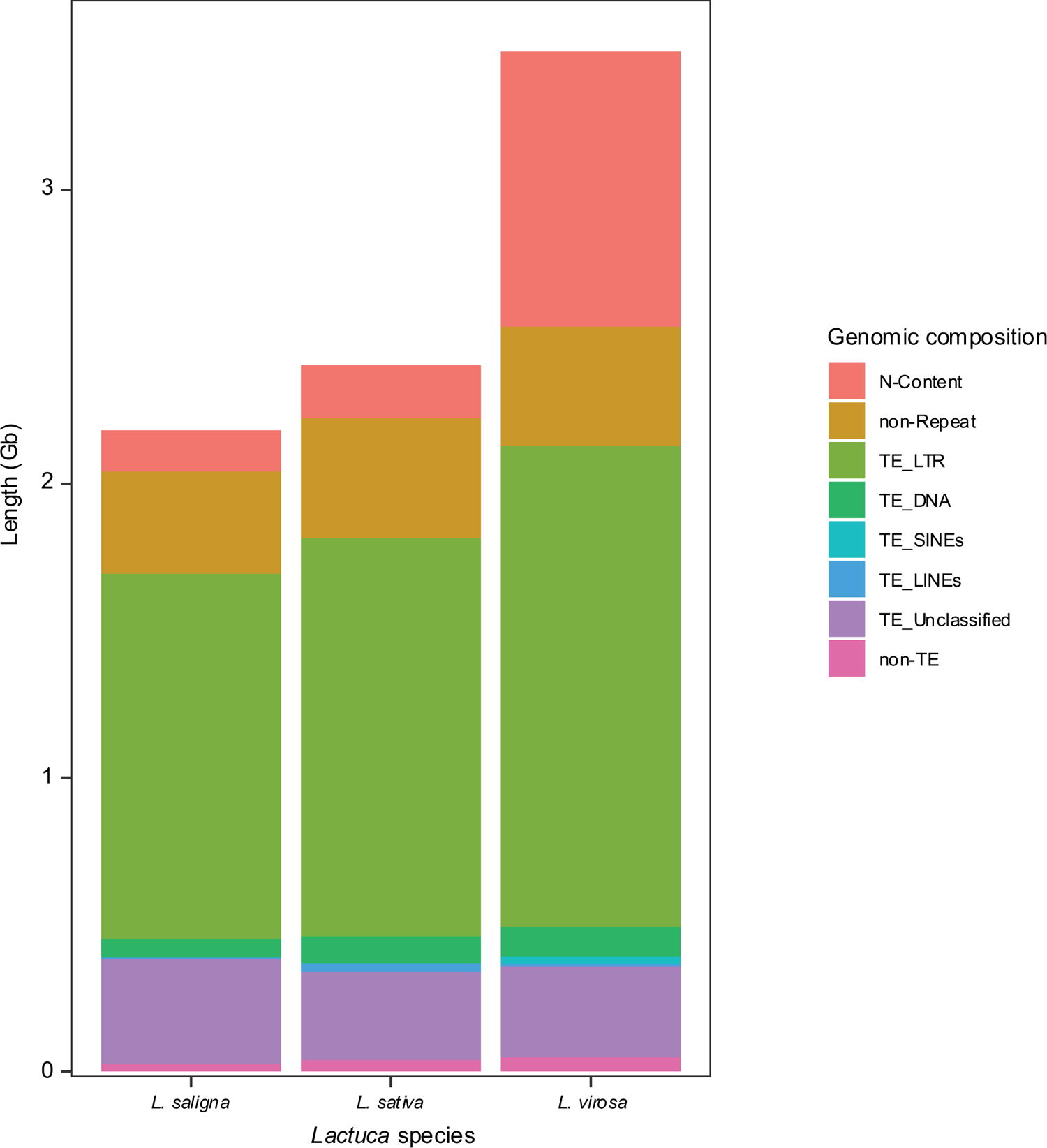
Genome composition of three *Lactuca* spp. Assemblies including and excluding N content (Supported by Supplementary Table 2).

**Supplementary Figure 6.**
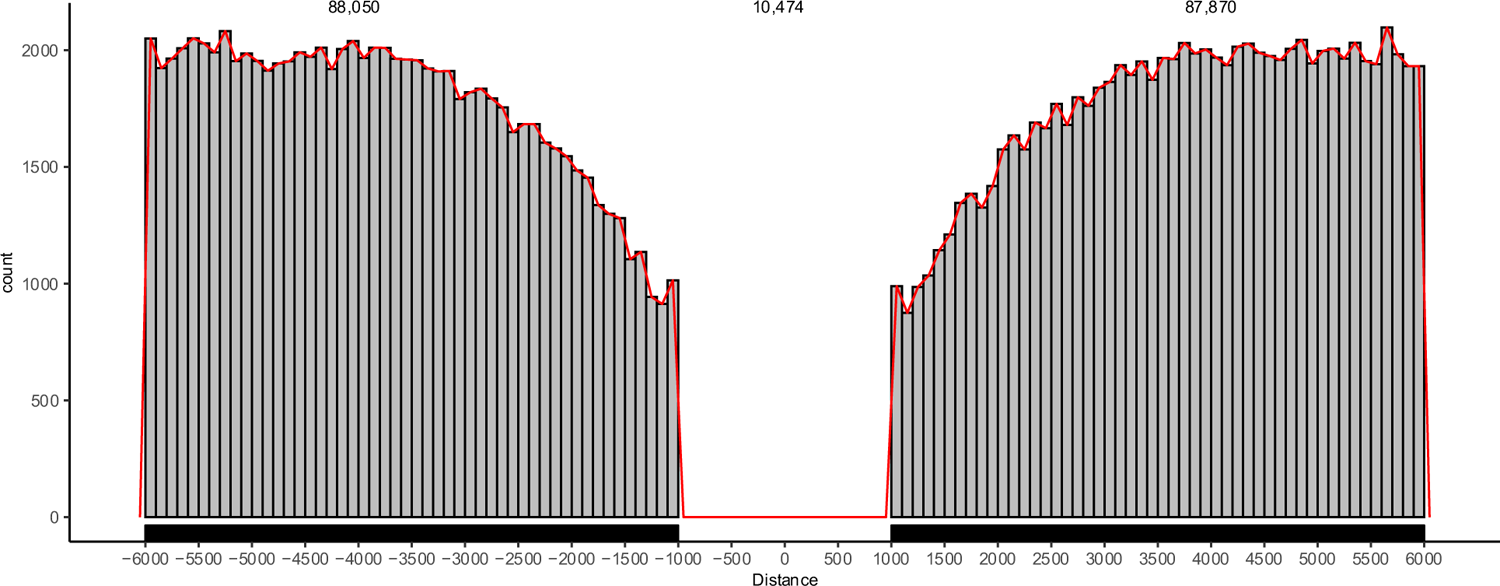
LTR distribution at gene flanking regions (+/- 5 kb) in the *L. virosa* genome. LTR extracted from the cross_match summary (.out) file of RepeatMasker. Data were compared to the genome using bedmap to determine whether an LTR is genic or intergenic. In total, there are 10,474 genic and 1,690,664 intergenic LTRs. Among intergenic LTRs, 88,050 and 87,870 are located at up and down 5 kb regions, respectively. This bar plot shows the density of LTR for the up and down 5 kb intergenic regions (bin = 100 bp).

**Supplementary Figure 7.**
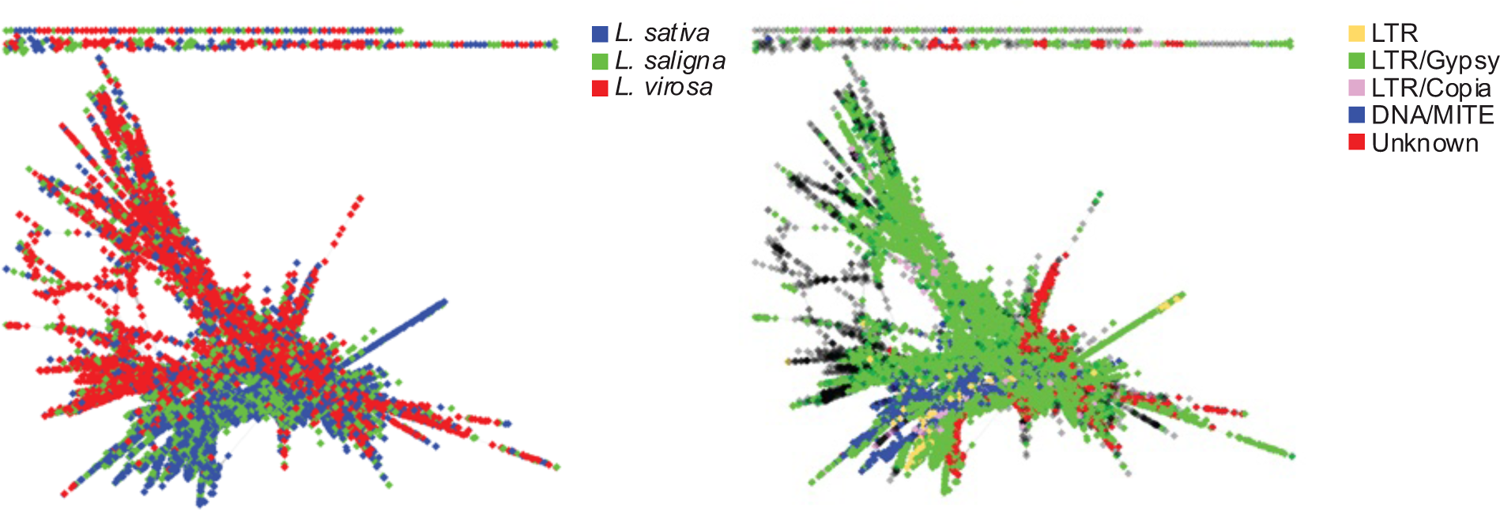
Example cluster of repeat reads from RepeatExplorer. An illustration of cluster 10 (Supercluster 2) from two perspectives. The left graph shows the read contribution from three species. The right cluster demonstrates the characterized transposable elements (TEs) in this cluster.

**Supplementary Figure 8.**
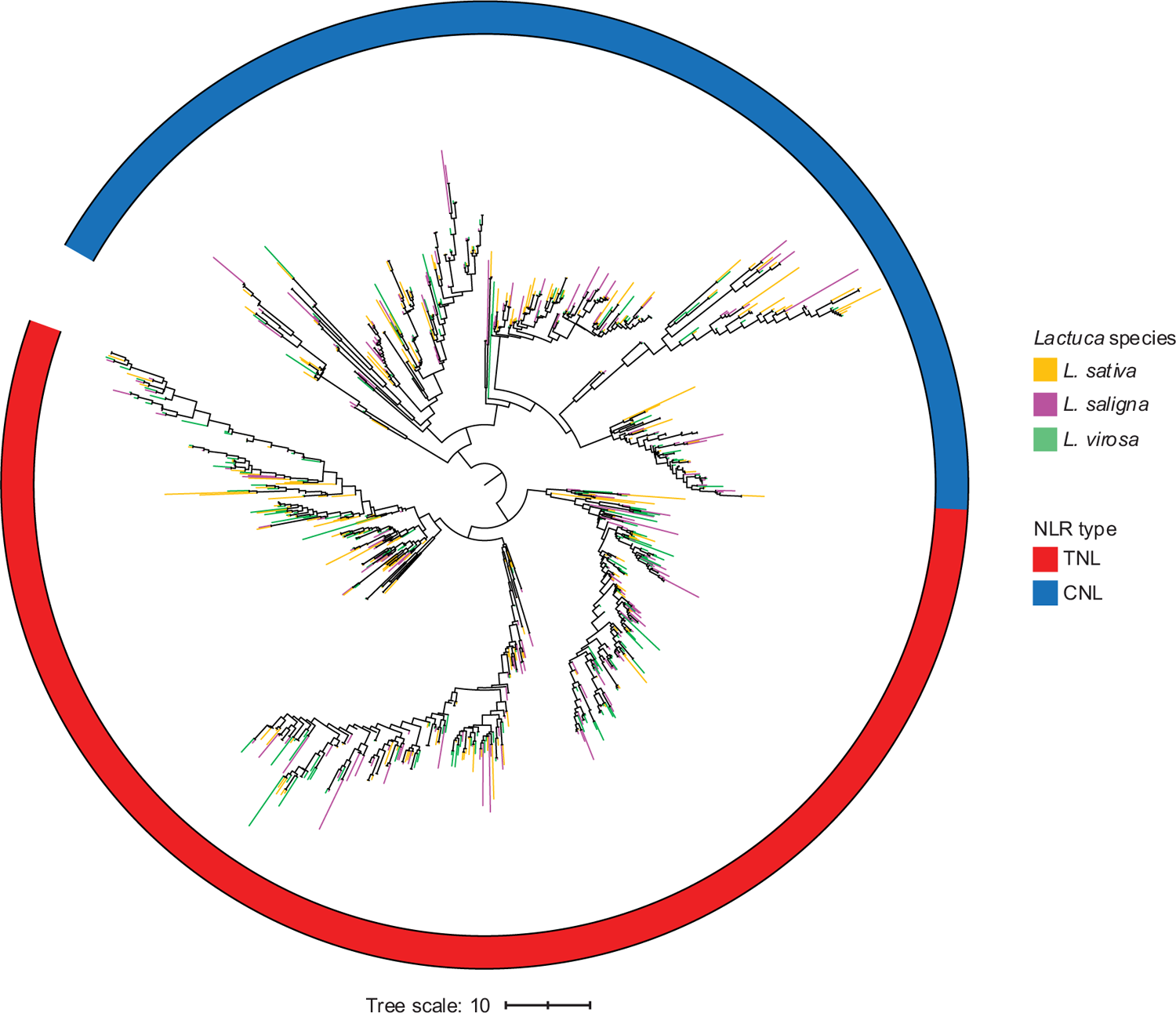
Circular tree of NLRs for *Lactuca* species generated by IQTREE. The tree was re-rooted at the midpoint between the TNL (NLR with TOLL/interleukin-1 receptor domain) and CNL (with coiled-coil domain) clade, and ultra-fast bootstrap approximation (UFBoot) support values were calculated with 1,000 repetitions. Branch color represents the three species: yellow for *L. sativa*, purple for *L. saligna*, and green for *L. virosa*.

## Supplementary Tables

**Supplementary Table 1.** Summary statistics of DNA and RNA sequencing.

| Type | Tissue | Technology | # reads/molecules | # bases |
| --- | --- | --- | --- | --- |
| DNA | Leaf | PacBio | 568,577 | 73,962,385,495 |
| DNA | Leaf | Illumina (raw) | 2,042,023,560 | 256,256,920,629 |
| DNA | Leaf | Illumina (Trimmed)* | 1,589,767,384 | 182,804,246,562 |
| DNA | Leaf | Hi-C | 898,000,000 | 134,700,000,000 |
| DNA | Leaf | Bionano | 5,199,031 | 684,445,635,000 |
| RNA | Root | Illumina | 182,472,598 | 22,594,657,038 |
| RNA | Leaf | Illumina | 178,858,548 | 22,182,587,010 |
| RNA | Flower | Illumina | 154,864,274 | 19,196,608,198 |

\*Trimmed Illumina reads used for genome size estimation by GenomeScope.

**Supplementary Table 2.** Overview of all statistics of *L. virosa* during the process of improving the assembly. N90 and L90 statistics are calculated for the scaffolded assembly.

| Step | # seq | Total length | N90 | L90 | Longest | Shortest | # Ns | # Gaps |
| --- | --- | --- | --- | --- | --- | --- | --- | --- |
| 00 input genome | 29 | 3305115164 | 278534532 | 9 | 442430716 | 90711 | 905096689 | 202872 |
| 01 polish | 29 | 3307727908 | 278766989 | 9 | 442776370 | 90711 | 904557320 | 116360 |
| 02 combine & purge | 54814 | 3569450220 | 76965895 | 11 | 442776370 | 1001 | 904854863 | 152711 |
| 03 filter non-nuclear | 54786 | 3569358957 | 76965895 | 11 | 442776370 | 1001 | 904854281 | 152696 |
| 04 filter contamination | 39955 | 3529432463 | 116478781 | 10 | 442776370 | 1001 | 904628957 | 146211 |
| 05 polish new | 39955 | 3534064810 | 116478781 | 10 | 442776370 | 930 | 903780369 | 128483 |
| 07 scaffold | 24767 | 3596785038 | 76965895 | 11 | 442776370 | 930 | 965886005 | 143368 |
| 08 filter RNA-seq | 6727 | 3449425426 | 116478781 | 10 | 442776370 | 1003 | 937474839 | 130693 |
| 09 filter length | 5855 | 3446681275 | 116478781 | 10 | 442776370 | 5010 | 937453665 | 130351 |

**Supplementary Table 3.** Summary of the different *L. virosa* assemblies used in this study.

| Assembly version | Initial assembly | Wei et al. (2021) assembly | Final assembly |
| --- | --- | --- | --- |
| Genome size | 3,305,115,164 | 2,912,862,072 | 3,446,681,275 |
| # seq | 29 | 3,694,810 | 5,855 |
| N50 scaffold | 316,657,893 | 4,910 | 316,905,932 |
| L50 scaffold | 5 | 137,357 | 5 |
| N90 scaffold | 278,534,532 | 201 | 116,478,781 |
| L90 scaffold | 9 | 1,468,356 | 10 |
| Assembly complete BUSCO | 75.20% | 92.70% | 96.20% |
| # genes | - | - | 39,887 |
| # transcripts | - | - | 42,791 |

**Supplementary Table 4.** Functional annotation summary*.

| Database | Number of matched genes | Percent of total genes (%) |
| --- | --- | --- |
| Interproscan | 31804 | 79.7 |
| SwissProt | 24362 | 61.1 |
| TrEMBL | 36941 | 92.6 |
| KEGG | 12853 | 32.2 |
| Combined | 37106 | 93.0 |

\*Supported by Supplementary Data 2A-B.

**Supplementary Table 5.** Tentative linkage groups (LG) of *L. virosa* scaffolds.

| Syntenic<br>lettuce LG | Original ID | Short ID | Length | Orientation |
| --- | --- | --- | --- | --- |
| Chr1 | Lvir_CGN04683_V4_scf6 | Scaffold6 | 313,930,088 | - |
| Chr1 | Lvir_CGN04683_V4_scf12 | Scaffold12 | 12,950,415 | - |
| Chr2 | Lvir_CGN04683_V4_scf5 | Scaffold5 | 316,905,932 | + |
| Chr3 | Lvir_CGN04683_V4_scf1 | Scaffold1 | 442,776,370 | - |
| Chr4 | Lvir_CGN04683_V4_scf2 | Scaffold2 | 416,659,169 | - |
| Chr4 | Lvir_CGN04683_V4_scf10 | Scaffold10 | 116,478,781 | - |
| Chr5 | Lvir_CGN04683_V4_scf3 | Scaffold3 | 385,159,916 | - |
| Chr5 | Lvir_CGN04683_V4_scf11 | Scaffold11 | 76,965,895 | - |
| Chr6 | Lvir_CGN04683_V4_scf9 | Scaffold9 | 278,766,989 | + |
| Chr7 | Lvir_CGN04683_V4_scf8 | Scaffold8 | 290,218,532 | + |
| Chr8 | Lvir_CGN04683_V4_scf4 | Scaffold4 | 342,977,835 | + |
| Chr9 | Lvir_CGN04683_V4_scf7 | Scaffold7 | 307,405,596 | - |

**Supplementary Table 6.** Percentage of sequence for different types of repeat elements in *Lactuca* spp. genomes (Supported by Supplementary Data 3A). For each species, both total percentage (Total) and percentage for the genome disregarding unknown bases (Excl N/X) are given.

| Species | *L. sativa* |  | *L. saligna* |  | *L. virosa* |  |
| --- | --- | --- | --- | --- | --- | --- |
| Classification | Total | Excl N/X** | Total* | Excl N/X** | Total* | Excl N/X** |
| SINEs | 0.03 | 0.03 | 0.01 | 0.01 | 0.72 | 0.99 |
| LINEs | 1.24 | 1.34 | 0.32 | 0.34 | 0.31 | 0.42 |
| LTR elements | 56.73 | 61.39 | 57.28 | 61.24 | 47.51 | 65.26 |
| DNA elements | 3.74 | 4.04 | 2.97 | 3.17 | 2.88 | 3.96 |
| Unclassified | 12.53 | 13.56 | 16.42 | 17.55 | 8.92 | 12.25 |
| Total TEs | 74.27 | 80.37 | 76.99 | 82.32 | 60.34 | 82.88 |
| Small RNA: | 0.03 | 0.04 | 0.17 | 0.19 | 0.17 | 0.24 |
| Satellites | 0.79 | 0.86 | 0.86 | 0.92 | 1.16 | 1.59 |
| Simple repeats | 0.78 | 0.85 | 0.12 | 0.13 | 0.07 | 0.09 |
| Low complexity | 0.00 | 0.00 | 0 | 0 | 0 | 0 |

\* Percentage of annotated repeat sequence calculated based on total genome length.
\*\* Percentage of annotated repeat sequence calculated based on genome length excluding N/X content.

**Supplementary Table 7.** Summary of individual and comparative RepeatExplorer analysis. Lsat is *L. sativa*, Lsal is *L. saligna* and Lvir is *L. virosa*.

| Analysis | Individual |  |  | Comparison |
| --- | --- | --- | --- | --- |
| Sample | Lsat | Lsal | Lvir | Lsat + Lsal + Lvir |
| Coverage of sampled reads (x) | 0.2 | 0.2 | 0.2 | 0.07 |
| Number of input reads | 4,166,668 | 3,833,334 | 6,166,668 | 4,958,336 |
| Number of analyzed reads | 3,933,112 | 3,651,552 | 3,865,971 | 4,830,991** |
| Coverage of analyzed reads (x) | 0.19 | 0.19 | 0.13 | 0.07 |
| Number of singlets | 814,812 | 827,510 | 685,087 | 982,735 |
| Reads in top clusters (%) | 68 | 65 | 74 | 71 |
| Reads in all clusters (%) | 79 | 77 | 82 | 80 |
| # top-clusters* | 264 | 245 | 239 | 266 |
| # superclusters | 162 | 150 | 153 | 171 |

\* Cluster size > 0.01% analyzed reads.
\*\* SAL = 1,307,006, SAT = 1,420,966, and VIR = 2,103,018.

**Supplementary Table 8.**
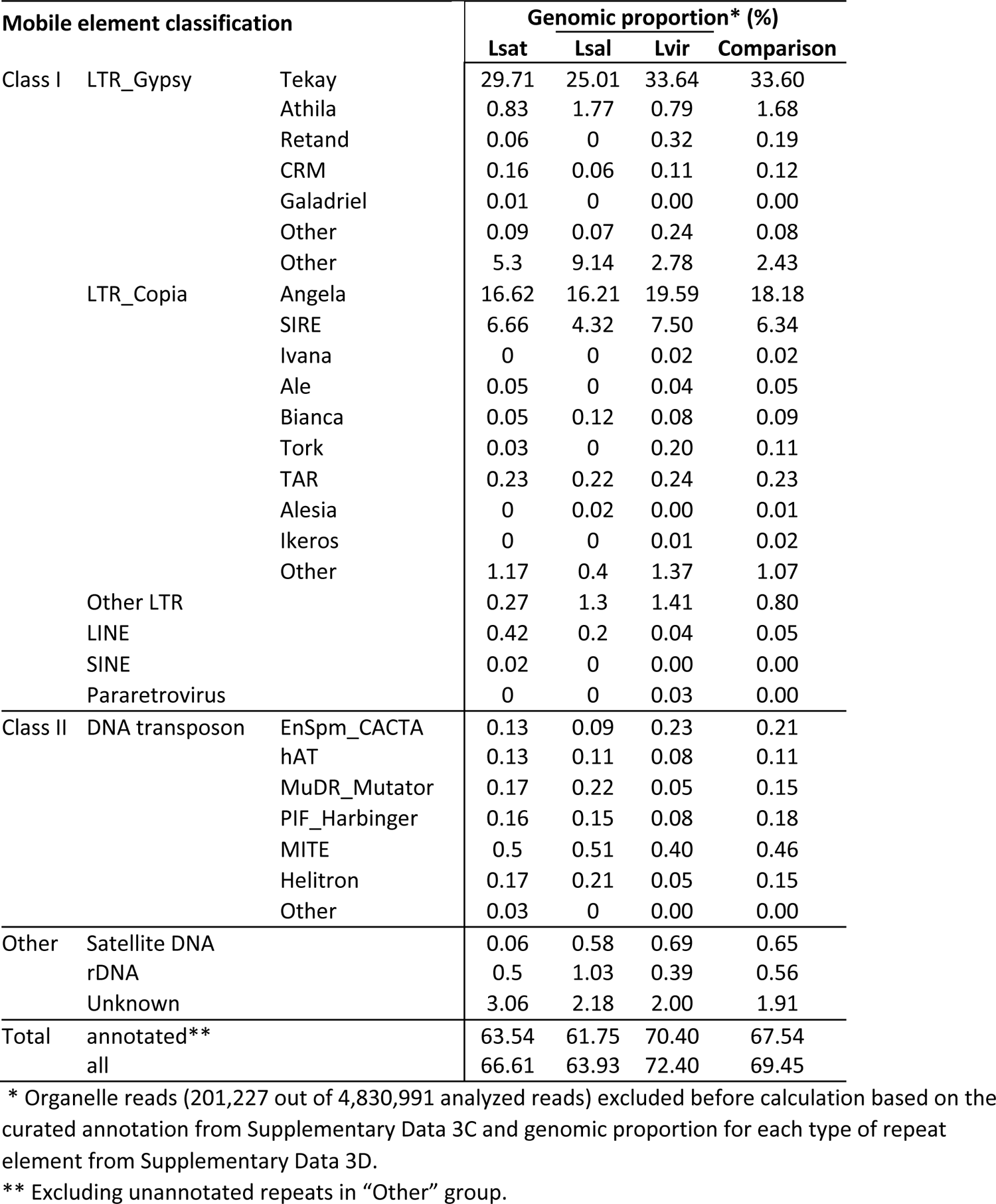
Genomic proportion of annotated clusters for individual and comparative analyses. Lsat is *L. sativa*, Lsal is *L. saligna* and Lvir is *L. virosa*.

**Supplementary Table 9.**
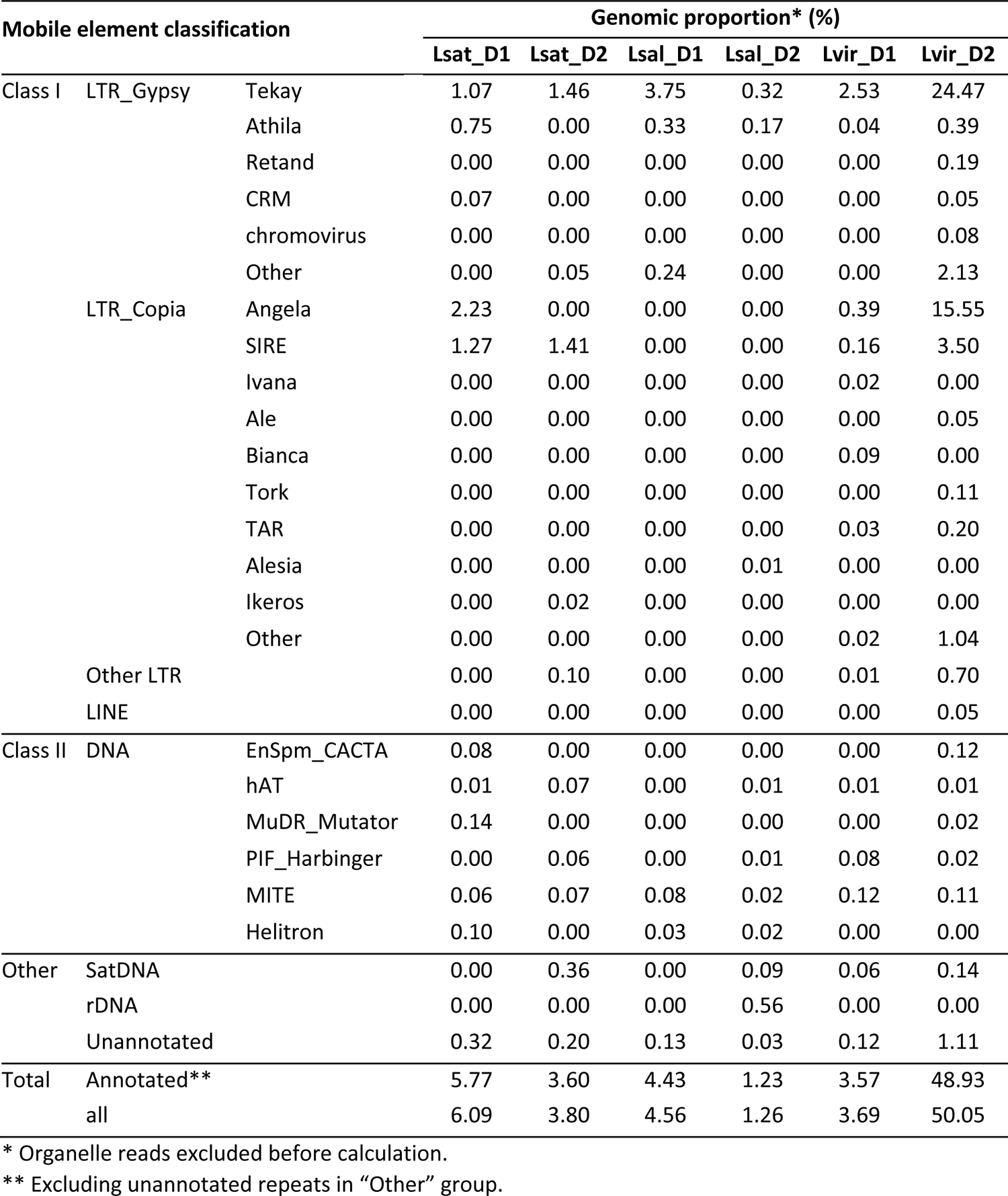
Genomic proportion of six groups after hierarchical clustering for annotated repeat clusters (supports Figure 4).

| Mobile element classification |  |  | Genomic proportion* (%) |  |  |  |  |  |
| --- | --- | --- | --- | --- | --- | --- | --- | --- |
|  |  |  | Lsat_D1 | Lsat_D2 | Lsal_D1 | Lsal_D2 | Lvir_D1 | Lvir_D2 |
| Class I | LTR_Gypsy | Tekay | 1.07 | 1.46 | 3.75 | 0.32 | 2.53 | 24.47 |
|  |  | Athila | 0.75 | 0.00 | 0.33 | 0.17 | 0.04 | 0.39 |
|  |  | Retand | 0.00 | 0.00 | 0.00 | 0.00 | 0.00 | 0.19 |
|  |  | CRM | 0.07 | 0.00 | 0.00 | 0.00 | 0.00 | 0.05 |
|  |  | chromovirus | 0.00 | 0.00 | 0.00 | 0.00 | 0.00 | 0.08 |
|  |  | Other | 0.00 | 0.05 | 0.24 | 0.00 | 0.00 | 2.13 |
|  | LTR_Copia | Angela | 2.23 | 0.00 | 0.00 | 0.00 | 0.39 | 15.55 |
|  |  | SIRE | 1.27 | 1.41 | 0.00 | 0.00 | 0.16 | 3.50 |
|  |  | Ivana | 0.00 | 0.00 | 0.00 | 0.00 | 0.02 | 0.00 |
|  |  | Ale | 0.00 | 0.00 | 0.00 | 0.00 | 0.00 | 0.05 |
|  |  | Bianca | 0.00 | 0.00 | 0.00 | 0.00 | 0.09 | 0.00 |
|  |  | Tork | 0.00 | 0.00 | 0.00 | 0.00 | 0.00 | 0.11 |
|  |  | TAR | 0.00 | 0.00 | 0.00 | 0.00 | 0.03 | 0.20 |
|  |  | Alesia | 0.00 | 0.00 | 0.00 | 0.01 | 0.00 | 0.00 |
|  |  | Ikeros | 0.00 | 0.02 | 0.00 | 0.00 | 0.00 | 0.00 |
|  |  | Other | 0.00 | 0.00 | 0.00 | 0.00 | 0.02 | 1.04 |
|  | Other LTR |  | 0.00 | 0.10 | 0.00 | 0.00 | 0.01 | 0.70 |
|  | LINE |  | 0.00 | 0.00 | 0.00 | 0.00 | 0.00 | 0.05 |
| Class II | DNA | EnSpm_CACTA | 0.08 | 0.00 | 0.00 | 0.00 | 0.00 | 0.12 |
|  |  | hAT | 0.01 | 0.07 | 0.00 | 0.01 | 0.01 | 0.01 |
|  |  | MuDR_Mutator | 0.14 | 0.00 | 0.00 | 0.00 | 0.00 | 0.02 |
|  |  | PIF_Harbinger | 0.00 | 0.06 | 0.00 | 0.01 | 0.08 | 0.02 |
|  |  | MITE | 0.06 | 0.07 | 0.08 | 0.02 | 0.12 | 0.11 |
|  |  | Helitron | 0.10 | 0.00 | 0.03 | 0.02 | 0.00 | 0.00 |
| Other | SatDNA |  | 0.00 | 0.36 | 0.00 | 0.09 | 0.06 | 0.14 |
|  | rDNA |  | 0.00 | 0.00 | 0.00 | 0.56 | 0.00 | 0.00 |
|  | Unannotated |  | 0.32 | 0.20 | 0.13 | 0.03 | 0.12 | 1.11 |
| Total | Annotated** |  | 5.77 | 3.60 | 4.43 | 1.23 | 3.57 | 48.93 |
|  | all |  | 6.09 | 3.80 | 4.56 | 1.26 | 3.69 | 50.05 |

\* Organelle reads excluded before calculation.
\*\* Excluding unannotated repeats in "Other" group.

**Supplementary Table 10.** Summary of NLR domain search for three *Lactuca* spp. (Supports Table 2).

| Type | Structure** | Number of gene |  |  |
| --- | --- | --- | --- | --- |
|  |  | *L. sativa* | *L. saligna* | *L. virosa* |
| CNL type* | CNL | 72 | 72 | 74 |
|  | CN | 4 | 11 | 12 |
|  | NcL | 71 | 34 | 48 |
|  | Nc | 11 | 22 | 14 |
|  | Total | 158 | 139 | 148 |
| TNL type | TNL | 184 | 133 | 106 |
|  | TN | 1 | 4 | 0 |
|  | NtL | 42 | 38 | 54 |
|  | Nt | 0 | 9 | 1 |
|  | Total | 227 | 184 | 161 |
| Total NLR |  | 385 | 323 | 309 |

\* RPW8 and Rx\_N type of CNL included in this study.
\*\* Capital letters represent the domain identified by HMMER search, while lowercase letters represent the NLR classification by phylogeny based on nucleotide-binding domain alignment.

**Supplementary Table 11.**
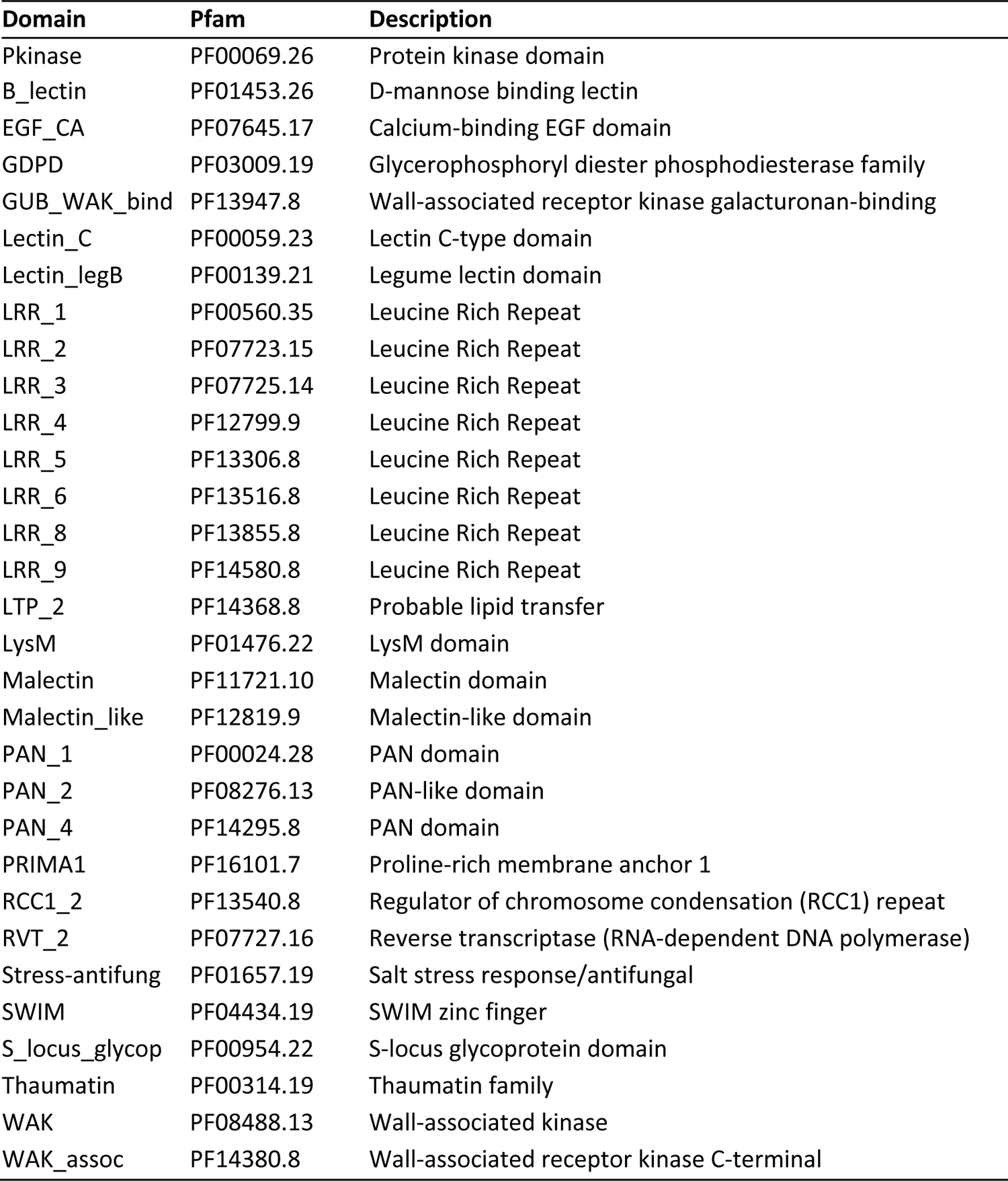
Pfam HMM motifs used for RLK classification (Supports Table 2).

**Supplementary Table 12.** RLK classification based on extracellular domain (Supports Table 2).

| Domain | Classification* |  |  |  |  |  |  |  |  |  |
| --- | --- | --- | --- | --- | --- | --- | --- | --- | --- | --- |
|  | Rcc1-RK | WAK | G-LecRK | L-LecRK | C-LecRK | CRK | LysM-RK | Malectin-RK | LRR-RK | PERK |
| RCC1_2 | x |  |  |  |  |  |  |  |  |  |
| GUB_WAK_bind |  | x |  |  |  |  |  |  |  |  |
| EGF_CA |  | x |  |  |  |  |  |  |  |  |
| WAK |  | x |  |  |  |  |  |  |  |  |
| WAK_assoc |  | x |  |  |  |  |  |  |  |  |
| B_lectin |  |  | x |  |  |  |  |  |  |  |
| PAN_2 |  |  | x |  |  |  |  |  |  |  |
| PAN_1 |  |  | x |  |  |  |  |  |  |  |
| PAN_4 |  |  | x |  |  |  |  |  |  |  |
| S_locus_glycop |  |  | x |  |  |  |  |  |  |  |
| Lectin_legB |  |  |  | x |  |  |  |  |  |  |
| Lectin_C |  |  |  |  | x |  |  |  |  |  |
| Stress-antifung |  |  |  |  |  | x |  |  |  |  |
| LysM |  |  |  |  |  |  | x |  |  |  |
| LRR_1 |  |  |  |  |  |  |  | x | x |  |
| LRR_2 |  |  |  |  |  |  |  | x | x |  |
| LRR_4 |  |  |  |  |  |  |  | x | x |  |
| LRR_5 |  |  |  |  |  |  |  | x | x |  |
| LRR_6 |  |  |  |  |  |  |  | x | x |  |
| LRR_8 |  |  |  |  |  |  |  | x | x |  |
| LRR_9 |  |  |  |  |  |  |  | x | x |  |
| LRR |  |  |  |  |  |  |  | x | x |  |
| Malectin |  |  |  |  |  |  |  | x |  |  |
| Malectin_like |  |  |  |  |  |  |  | x |  |  |
| PRIMA1 |  |  |  |  |  |  |  |  |  | x |

\* Abbreviation for classified RLKs:
RCC1-RK RCC1 repeat receptor-like kinase
WAK Wall-associated receptor-like kinase
G-LecRK G-type lectin receptor kinase
L-LecRK L-type lectin receptor kinase
C-LecRK C-type lectin receptor kinase
CRK cysteine-rich receptor-like kinases
LysM-RK Lysin motif receptor-like kinase
Malectin-RK Malectin receptor-like kinase

## Supplementary Data

Datasets are available in: https://figshare.com/s/aa3ec2f08495f0d24dfb

**Supplementary Data 1**. Genome assembly and scaffolding. **A**, The input assembly scaffolds in the Hirise scaffolds using Hi-C. **B**, The input assembly scaffolds in the final scaffolds (WUR + BGI). **C**, The sequence length of scaffolds in final assembly (WUR + BGI).

**Supplementary Data 2**. Functional annotation and homology grouping of *L. virosa* transcripts. **A**, Detailed match of functional annotation by different approaches for *L. virosa* transcripts for all isoforms. **B**. Detailed match of functional annotation by different approaches for *L. virosa* genes. **C**, Homology groups of the three *Lactuca* species calculated by PanTools. **D**, InterPro enrichment of *L. virosa* specific homologs.

**Supplementary Data 3**. Repeatome analysis of *L. virosa*, *L. sativa* and *L. saligna*. **A**, Summary of RepeatMasker output for the three genomes. **B**, Matrix and cumulative stats of RepeatExplorer clusters. **C**, Curated annotation of RepeatExplorer clusters. **D,** Genomic proportion of curated clusters excluding organelle reads.

**Supplementary Data 4**. Identified NLR and RLK proteins in *L. virosa* and *L. saligna*. **A**, Identification and classification of NLR (*L. virosa*). **B**, Identification and classification of NLR (*L. sativa*). **C**, Identification and classification of RLK (*L. virosa*). **D**, Identification and classification of RLK (*L. sativa*).

**Supplementary Data 5**. Overview of homology within NLRs and RLKs in *Lactuca*. **A**, Homology groups of identified NLRs and RLKs for three *Lactuca* species. **B**, Tandem array detection of identified NLRs and RLKs for three *Lactuca* species.

